# Typical and disrupted small-world architecture and regional communication in full-term and preterm infants

**DOI:** 10.1101/2023.06.12.544595

**Authors:** Huiqing Hu, Peter Coppola, Emmanuel Stamatakis, Lorina Naci

## Abstract

One fundamental property of conscious experiences is that they are both differentiated and integrated. Adult functional brain networks exhibit an elegant “small-world” architecture. This optimal architecture enables efficient and cost-effective localized information processing and information integration between long-distance regions across the brain. It remains unclear whether the functional small-world architecture is developed in neonates at birth and how this development may be altered by premature birth. To address this gap, we investigated the development of small-world architecture in neonates. To understand the effect of early neonate age on small-world architecture, we also assessed neonates born prematurely or before term-equivalent age (TEA). We used the Developing Human Connectome Project (dHCP), a large neonatal functional magnetic resonance imaging (MRI) dataset with high temporal and spatial resolution. Resting-state functional MRI data for full-term neonates (N = 278, age 41.2 weeks ± 12.2 days) and preterm neonates scanned at TEA (N = 72, 40.9 weeks ± 14.6 days), or before TEA (N = 70, 34.7 weeks ± 12.7 days), were obtained from the dHCP, and for a reference adult group (N = 176, 22–36 years), from the Human Connectome Project. Whole-brain functional network properties were evaluated with comprehensive spatial resolution using graph theoretical analyses. Although different from the adults’, small-world architecture was developed in full-term born neonates at birth. Premature neonates before TEA showed dramatic underdevelopment of small-world architecture and regional communication in 9/11 brain networks, with disruption in 32% of nodes primarily distributed within the somatomotor, dorsal attention, cingulo-opercular, and frontoparietal control network. By TEA, premature neonates showed large-scale recuperation of regional communication, with 1.4% of nodes, distributed in the frontoparietal, salience, and visual networks remaining significantly underdeveloped. Our results suggest that, at full-term birth or by term-equivalent age, infants possess well-developed small-world architecture, which facilitates differentiated and integrated neural processes that give rise to conscious experiences. Conversely, they suggest that this brain infrastructure is significantly underdeveloped before infants reach term-equivalent age. These findings improve understanding of the ontogeny of functional small-world architecture and efficiency of neural communication across distinct brain networks in infants at birth.

## Introduction

One fundamental property of conscious experiences is that they are both differentiated and integrated (Deco et al., 2015; Seth, 2009; Seth, et al., 2006; Cooney & Gazzaniga, 2003; Tononi & Edelman, 1998). A conscious brain can differentiate and integrate different experiential features (e.g., sights, sounds, smells, visceral sensations, etc.) into a unified conscious experience. It has been postulated that the balance of differentiating and integrative processes, which appear prima facia to be opposed, is a key feature of consciousness (Seth et al., 2006). Graph theoretical analyses of brain activity have shown that functional brain networks display a so-called ‘small-world’ organization, comprised of dense within-network connections that facilitate information specialization and sparse between-network connections that enhance global information integration (Bassett & Bullmore, 2017; 2006).

Small-world brain organization is, thus, an important feature that facilitates the optimal balance of information segregation and integration required for conscious processing. Studies found that small-world organization is modulated by consciousness state manipulations (i.e., by sleep or anaesthesia) (Luppi et al., 2021; Barttfeld et al., 2015; Uehara et al., 2014; Monti et al., 2013; Schröter et al., 2012), and by consciousness perturbation after brain injury (Duclos et al., 2021; Luppi et al., 2019), further providing empirical evidence that small-world organization supports conscious information processing. For example, Luppi et al. (2019) observed that patients with disorders of consciousness (i.e., in the unresponsive wakefulness syndrome/vegetative state, or the minimally conscious state) displayed reduced small-world characteristics relative to awake healthy volunteers. Other studies have also shown similar effects with decreasing level of consciousness, for example, a reduction of long- and short-range functional brain connections in anaesthesia (Schröter et al., 2012), or reduction of long-range functional communication in sleep (Uehara et al., 2014).

There is broad consensus from both theoretical and empirical studies that small-word organization is central to processes that support human consciousness in adults (see Monti et al., (2013); Schröter et al., (2012) for a divergent view). However, it remains poorly understood whether functional small-world architecture is present in neonates and how premature birth affects it. A literature search in PubMed with keywords ‘small-world architecture’ or ‘small worldness’ or ‘small-world propensity’, ‘infant’ or ‘newborn’ or ‘neonate’ and ‘fMRI’ or ‘functional imaging,’ revealed inconsistent findings (Table 1). Some previous studies have suggested that the small-world architecture is present in full-term born neonates (Bouyssi-Kobar et al., 2019; Asis-Cruz et al., 2015; Omidvarni et al., 2014; Fransson et al., 2011), but several limitations undermine these studies’ conclusions. The index (σ) (Humphries & Gurney, 2008) used to quantify small-world properties in these studies has been criticised (Bassett & Bullmore, 2017; Papo et al., 2016; Telesford et al., 2011) because it can result in a high false-positive rate (Telesford et al., 2011). In addition, the small sample size (N = 10 to 66) and low temporal and spatial resolution limit the result reliability in these studies.

**Table 1.**
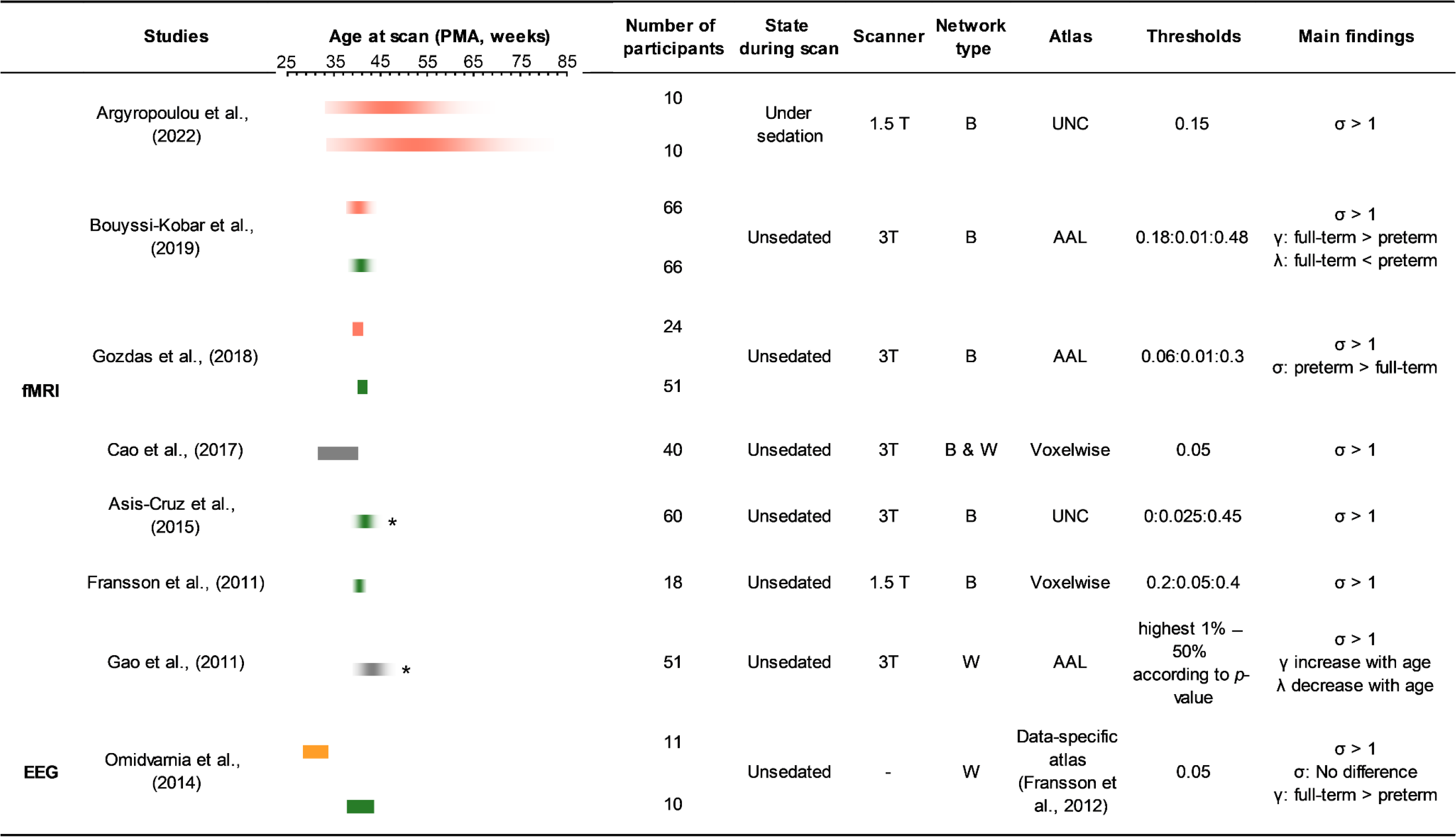
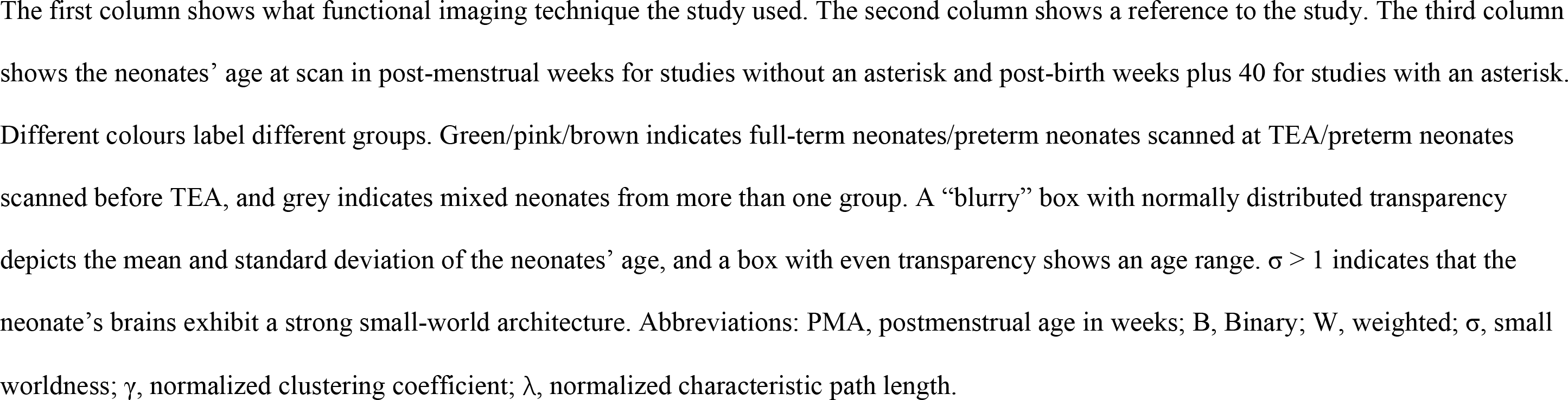
A summary of functional imaging studies that investigated the development of small-world architecture in neonates.

|  | Studies | Age at scan (PMA, weeks) |  |  |  |  |  |  | Number of participants | State during scan | Scanner | Network type | Atlas | Thresholds | Main findings |
| --- | --- | --- | --- | --- | --- | --- | --- | --- | --- | --- | --- | --- | --- | --- | --- |
|  |  | 25 | 35 | 45 | 55 | 65 | 75 | 85 |  |  |  |  |  |  |  |
| fMRI | Argyropoulou et al., (2022) | | | | | | | | 10 | Under sedation | 1.5 T | B | UNC | 0.15 | $\sigma > 1$ |
|  |  |  |  |  |  |  |  |  | 10 |  |  |  |  |  |  |
| | Bouyssi-Kobar et al., (2019) | | | | | | | | 66 | Unsedated | 3T | B | AAL | 0.18:0.01:0.48 | $\sigma > 1$<br>$\gamma$: full-term > preterm<br>$\lambda$: full-term < preterm |
|  |  |  |  |  |  |  |  |  | 66 |  |  |  |  |  |  |
| | Gozdas et al., (2018) | | | | | | | | 24 | Unsedated | 3T | B | AAL | 0.06:0.01:0.3 | $\sigma > 1$<br>$\sigma$: preterm > full-term |
|  |  |  |  |  |  |  |  |  | 51 |  |  |  |  |  |  |
| | Cao et al., (2017) | | | | | | | | 40 | Unsedated | 3T | B & W | Voxelwise | 0.05 | $\sigma > 1$ |
| | Asis-Cruz et al., (2015) | | | | | | | | 60 | Unsedated | 3T | B | UNC | 0:0.025:0.45 | $\sigma > 1$ |
| | Fransson et al., (2011) | | | | | | | | 18 | Unsedated | 1.5 T | B | Voxelwise | 0.2:0.05:0.4 | $\sigma > 1$ |
| | Gao et al., (2011) | | | | | | | | 51 | Unsedated | 3T | W | AAL | highest 1% – 50% according to p-value | $\sigma > 1$<br>$\gamma$ increase with age<br>$\lambda$ decrease with age |
| EEG | Omidvarnia et al., (2014) | | | | | | | | 11 | Unsedated | - | W | Data-specific atlas (Fransson et al., 2012) | 0.05 | $\sigma > 1$<br>$\sigma$: No difference<br>$\gamma$: full-term > preterm |
|  |  |  |  |  |  |  |  |  | 10 |  |  |  |  |  |  |

The first column shows what functional imaging technique the study used. The second column shows a reference to the study. The third column shows the neonates' age at scan in post-menstrual weeks for studies without an asterisk and post-birth weeks plus 40 for studies with an asterisk. Different colours label different groups. Green/pink/brown indicates full-term neonates/preterm neonates scanned at TEA/preterm neonates scanned before TEA, and grey indicates mixed neonates from more than one group. A “blurry” box with normally distributed transparency depicts the mean and standard deviation of the neonates' age, and a box with even transparency shows an age range. $\sigma > 1$ indicates that the neonate's brains exhibit a strong small-world architecture. Abbreviations: PMA, postmenstrual age in weeks; B, Binary; W, weighted; $\sigma$ , small worldness; $\gamma$ , normalized clustering coefficient; $\lambda$ , normalized characteristic path length.

To the best of our knowledge, only three small studies have examined small-world organization in neonates born at full term or term-equivalent age (TEA, postmenstrual age of 37 weeks) (Bouyssi-Kobar et el., 2019; Gozdas et al., 2018; Omidvarnia et al., 2014) and only two (Cao et al., 2017; Omidvarni et al., 2014) in neonates before TEA, in very small samples (N = 10 to 23). These studies have offered inconsistent results, likely due to different neuroimaging techniques, different brain atlases, and the small participants numbers. For example, Bouyssi-Kobar et al. (2019) used resting-state fMRI (rs-fMRI) and reported decreased clustering coefficient and increased path length in neonates at TEA (N = 66) relative to full-term born (N = 66), but no significant difference in small-worldness. Gozdas et al. (2018) used rs-fMRI and found significantly higher small worldness in preterm neonates (N = 24) compared with full-term born neonates (N = 51). Omidvarnia et al. (2014) used electroencephalography (EEG) and reported decreased clustering coefficient in preterm neonates before TEA (N = 11) but no significant difference in either small-worldness or path length to full-term born (N = 10).

Importantly, brain regions develop at different rates, and thus, the effects of premature birth on the balance of integration and segregation at the regional level in neonates remain unknown. For example, the sensorimotor network rapidly develops from around 31 to 41 weeks of gestational age and has an adult-like topology by term/TEA (Truzzi & Cusack, 2022; Dall’Orso et al., 2022; Allievi et al., 2016; Thomason et al., 2015; Doria et al., 2010; Smyser et al., 2010), while the visual network is underdeveloped at birth, and matures gradually in the first year of life (Truzzi & Cusack, 2022; Lemaître et al., 2021; Gao et al., 2015). Furthermore, before TEA, the dorsal attention network is present in preterm neonates, while the default mode network and executive network are dramatically underdeveloped at this time (Hu et al., 2022). Premature birth, in and of itself, has also been associated with differential atypical development of functional architecture within and across brain networks (Fenn-Moltu et al., 2023; Hu et al., 2022; Allievi et al., 2016).

To address these gaps, in this study, we first investigated whether small-world organization was developed at birth. We then asked how the balance of segregation and integration across the whole brain, and at the regional level where affected by premature birth and neonate age. To address the limitations of previous studies, we used rs-fMRI data from the largest open-source neonate dataset, the Developing Human Connectome Project (dHCP), which conferred several advantages, including a large sample size (N = 278), 3T magnetic resonance imaging (MRI), multiband echo-planar imaging (EPI) that significantly improves temporal resolution and signal-to-noise relative to sequences used in previous studies (Zhang et al., 2019), registration to more accurate week-to-week neonate structural templates, and significant improvements in motion correction and signal-to-noise ratio relative to previous studies (see Methods for further details). We also included a large reference adult group (N = 176) from the Human Connectome Project. The effect of chronological age at the time of assessment was deconfounded from the effect of premature birth (Bhutta et al., 2002) by the inclusion of two groups: the first (N = 72) born before and scanned at TEA, and the second (N = 70) born and scanned before TEA. A subset of paired scans collected from the same infants (N = 33) both before, and at TEA, was also included. We reasoned that any differences between neonates born and scanned at full-term and preterm neonates scanned at TEA would reflect the effects of premature birth, while controlling for neonate age. Conversely, any differences between preterm neonates scanned at TEA and those scanned before TEA would reflect the effects of neonate age. The balance of segregation and integration across the brain was assessed with rigorous measures of small-world architecture (Muldoon et al., 2016), whereas at the regional level with measures of nodal efficiency (Achard & Bullmore, 2007). Based on previous work (Hu et al., 2022), we tested the hypotheses that relative to full-term birth, (a) premature birth is associated with altered small-world architecture and nodal efficiency, and (b) that some alterations in these measures would persist as preterm neonates reach TEA.

## Materials and methods

### Participants

*Neonates*. The neonate data were from the second (2019) developing human connectome project (dHCP) public data release (http://www.developingconnectome.org/second-data-release/). All neonates were scanned at the Evelina Newborn Imaging Centre, Evelina London Children’s Hospital. Ethical approval was obtained from the UK’s National Research Ethics Committee, and parental informed consent was obtained before imaging. We used 278 full-term neonates (gestational age (GA) at birth = 39.9 weeks ± 8.6 days; postmenstrual age (PMA) at scan = 41.2 weeks ± 12.2 days; 119 females) after quality control procedures. 72 scans from the preterm neonates at TEA (GA at birth = 32.1 weeks ± 25.5 days; PMA at scan = 40.9 weeks ± 14.6 days; 32 females) and 70 scans from the preterm neonates before TEA (GA at birth = 32.5 weeks ± 20.7 days; PMA at scan = 34.7 weeks ± 12.7 days; 22 females) were included after quality control. We also included a subset of paired scans collected from the same preterm infants (N = 33, GA at birth = 32.2 weeks ± 20.1 days) at (PMA at scan = 41.0 weeks ± 10.16 days) and before TEA (PMA at scan = 34.5 weeks ± 11.5 days) to investigate the effects of neonate age. Further details on inclusion criteria are shown in Figure 1 and Table S1.

**Figure 1.**
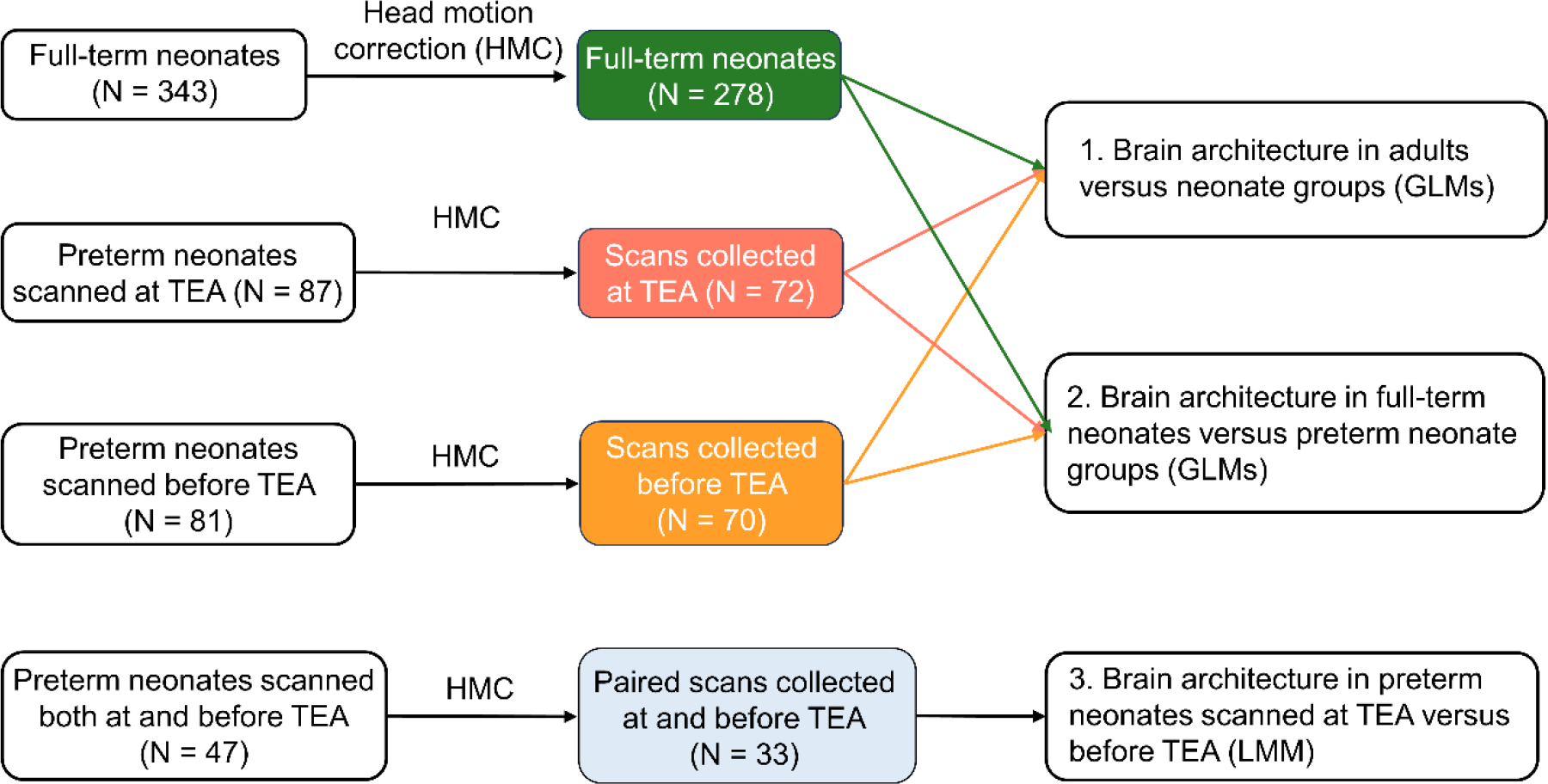
The number of scans included in data analyses. The frames in green/pink/brown indicate the scans collected from full-term neonates/preterm neonates at/before term-equivalent age that passed head motion criteria. The frame in light blue indicates the scans collected at and before term-equivalent age from the same preterm neonates. Abbreviations: TEA, term-equivalent age; GLMs, general linear models; LMM, linear mixed-effect model.

*Adults.* As a reference adult group, we used a subset (N = 176; 22 – 36 years; 99 females) of high-quality resting-state functional neuroimaging data from the final release of the Washington University-Minnesota Consortium of Human Connectome Project (HCP) selected by Ito et al. (2020). For details of study procedures see Van Essen et al. (2013).

### Data acquisition and Preprocessing

*dHCP*. Data were acquired on a 3T Philips Achieva with a dedicated neonatal imaging system including a neonatal 32-channel phased-array head coil. Fifteen minutes of high temporal and spatial resolution rs-fMRI data were acquired using a multislice gradient-echo echo planar imaging (EPI) sequence with multiband excitation (TE = 38 ms; TR = 392 ms; MB factor = 9x; 2.15 mm isotropic, 2300 volumes). In addition, single-band EPI reference (sbref) scans were also acquired with bandwidth-matched readout, along with additional spin echo EPI acquisitions with 4xAP and 4xPA phase-encoding directions. To correct susceptibility distortion in rs-fMRI data, field maps were also obtained from an interleaved (dual TE) spoiled gradient-echo sequence (TR = 10 ms; TE1 = 4.6 ms; TE2 = 6.9 ms; flip angle = 10°; 3 mm isotropic in-plane resolution, 6mm slice thickness). High-resolution T1- and T2-weighted anatomical imaging were also acquired in the same scan session, with a spatial resolution of 0.8 mm isotropic. For T1w image: TR = 4795 ms and the field of view (FOV) = 145 × 122 × 100 mm. For T2w image: TR = 12000 ms, TE = 156 ms and the FOV = 145 × 122 × 100 mm.

The dHCP rs-fMRI data were pre-processed by the dHCP group using the project’s in-house pipeline optimized for neonatal imaging (Fitzgibbon et al., 2020). In brief, motion and distortion correction and temporal high-pass filter procedures were applied. The cardiorespiratory fluctuations and multiband acquisition artefacts, the 24 extended rigid-body motion parameters together with single-subject ICA noise components were regressed out to reduce signal artefacts related to physiological noise and head movement (See SI methods for details). Then, we applied intensity normalization to the pre-processed dHCP rs-fMRI data to keep the preprocessing procedure consistent between neonates and adults. Last, a temporal low-pass filter (0.08 Hz low-pass cutoff) was applied to the neonatal data, as previous studies found that oscillations were primarily detected within grey matter in 0.01 – 0.08 Hz., (Zuo et al., 2010; Salvador et al., 2008).

To further reduce the effect of motion on graph theoretical measures, we applied a scrubbing procedure to retain a continuous sub-sample of the data (1600 time points, ∼70%) with the lowest motion for each neonate. See our previous paper (Hu et al., 2022) for details of the scrubbing procedure. Subjects with more than 160 motion-outlier volumes (10% of the cropped dataset) in the continuous subset were labelled ‘high level of motion’ and excluded entirely. Thus, 61 full-term neonates, 14 preterm neonates scanned at TEA, and 8 preterm neonates scanned before TEA were excluded. In addition, participants whose mean framewise displacement (FD) after the scrubbing procedure was larger than 0.5mm were also excluded. Thus, 4 full-term neonates, 1 preterm neonate scanned at TEA, and 3 preterm neonates scanned before TEA were excluded (Figure 1).

*HCP.* Data were acquired on a customized 3T Siemens “Connectome Skyra” with a 32- channel head coil. Resting-state images were collected using gradient-echo EPI sequence: TR = 720 ms; TE = 33.1 ms; flip angle = 52°; FOV = 208 × 180 mm (RO × PE), slice thickness = 2 mm, 72 slices, 2.0 mm isotropic voxels, 1200 volumes per run. The HCP rs-fMRI data were pre-processed by the HCP group (Glasser et al., 2013) using the following pipeline: distortion correction, motion correction, alignment of the original rs-fMRI data to the Montreal Neurological Institute (MNI) template space, intensity normalization, temporal high-pass filter, and denoising using ICA-FIX. The detailed pre-processing procedure can be found in SI Methods section. Then, we applied a temporal low-pass filter (0.08 Hz low-pass cutoff) to the pre-processed adult data.

As the selection of the subset of HCP had controlled head motion (i.e., exclusion of participants that had any fMRI run in which more than 50% of TRs had greater than 0.25mm FD), and adults generally have smaller maximal head motion than neonates, we did not apply the same scrubbing method used in the dHCP dataset to adult data. In addition, we randomly chose a continuous sub-sample of data time-series (871 time points) in adults as in neonates.

### Data analyses

#### Node definition

Using a meta-analytically derived functional brain atlas from Power at al., (2011), we defined 264 cortical and subcortical regions of interest (ROIs) (8-mm radius spheres) in MNI space. 33 uncertain nodes and 5 cerebral nodes were excluded and the remaining 226 nodes were from 11 networks: 1) sensory/somatomotor hand (30 nodes), 2) sensory/somatomotor mouth (5 nodes), 3) cingulo-opercular control (14 nodes), 4) auditory (13 nodes), 5) default mode (57 nodes), 6) visual (31 nodes); 7) frontoparietal control (25 nodes), 8) salience (18 nodes), 9) subcortical (13 nodes), 10, ventral attention (9 nodes), and dorsal attention (11 nodes) networks. For neonates, the 226 ROIs in MNI space were transformed into neonate native space using the following two-step procedure. We first aligned the ROIs in MNI space with the 40-week dHCP T1w template (Schuh et al., 2018) using the warp file generated from the transformation between the MNI T1w template and 40-week dHCP T1w template using ANTs (See SI Methods for details). Then, the nodes in 40-week dHCP T1w template space were transformed into neonate native space by applying the inverted func-to-template warp provided by the dHCP group (Fitzgibbon et al., 2020) using FSL. Time courses were extracted from each neonate based on these ROIs in native space and from adults based on ROIs in the MNI space.

To ensure that we used the same number of nodes/connections in the following graph construction, we checked the extracted time courses from adults, full-term neonates, preterm neonates at TEA, and preterm neonates before TEA. 9 nodes were further excluded because of low registration accuracy according to the following criteria: 1) over 5% of participants in any of the four groups missed time courses in that node; 2) node showed significant differences in the proportions of participants having missing time courses between the four groups. Thus, 4 nodes of DMN (node 83, 116, 119, 120), 4 nodes of the visual network (node 153, 165, 168, 172), and node 179 of the frontoparietal control network were excluded (Figure S10).

#### Graph construction

Pearson’s correlation between time courses of any two pairs of nodes was calculated. Then, self-connections were set to 0, and NaNs and negative correlations were removed. 21 proportional thresholds, from 10% to 30% in 1% increases (Godwin et al., 2015; Monti et al., 2013), were used to threshold functional matrices (Figure 2) to ensure that results were not driven by specific connection densities (Hallquist et al., 2018; van den Heuvel et al., 2017; Rubino & Sporns, 2010).

**Figure 2.**
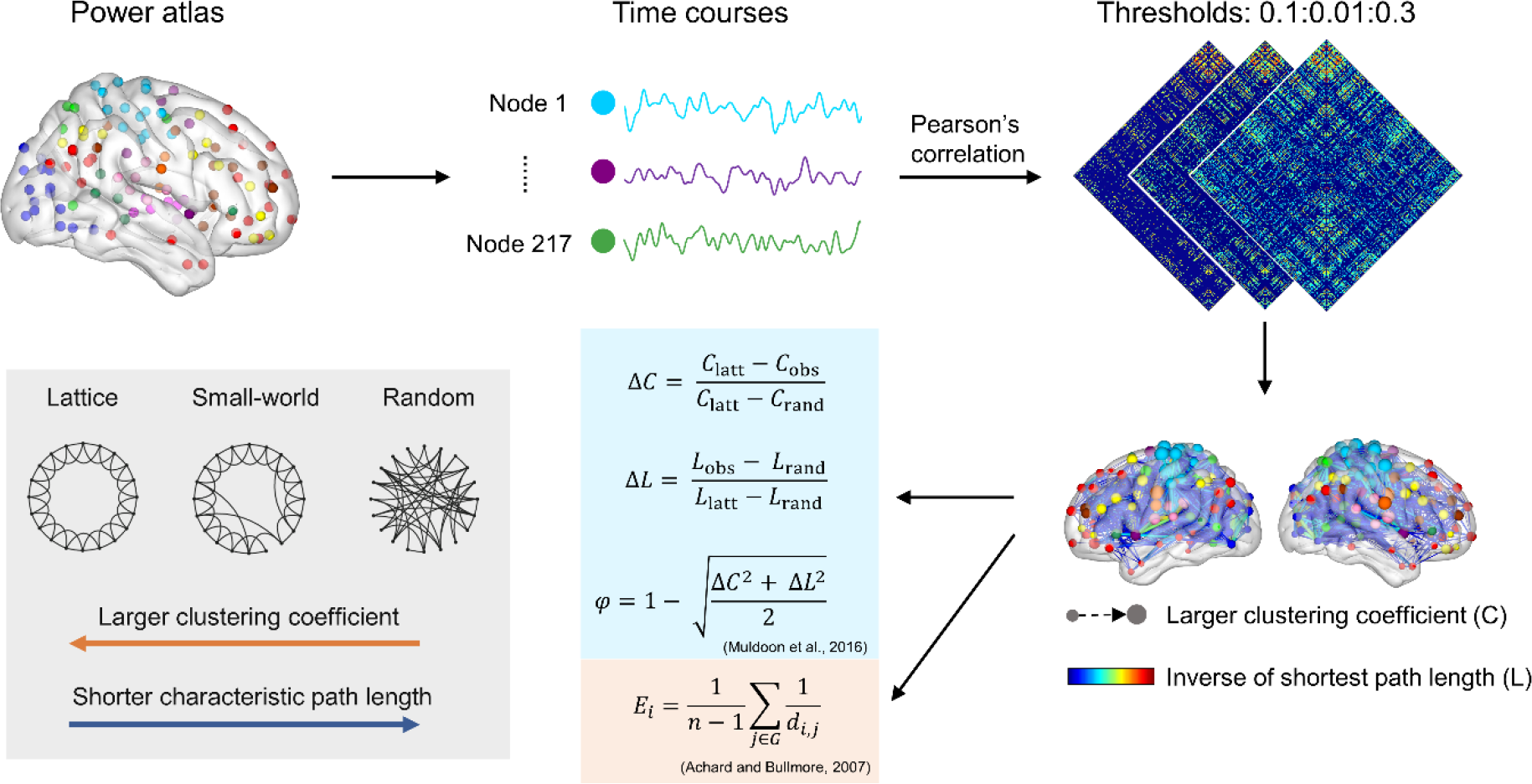
Schematic of the method. Time courses were extracted from each brain node, and Pearson’s correlation was applied to obtain a weighted graph for each participant. 21 proportional thresholds from 10% to 30% in 1% increases were applied to threshold the weighted graphs. The local clustering coefficient for each node and the shortest path length between every two nodes were calculated based on weighted graphs at each cost function. Nodal efficiency for each node was calculated based on shorted path length at each cost function and then averaged. Then, small-world propensity, normalized clustering coefficient and normalized characteristic path length for each brain network were calculated (Muldoon et al., 2016) at each cost function and averaged.

#### Small-world properties

Nodal-level clustering coefficient and shortest path length were first calculated based on the FC matrices under each threshold for each participant. The clustering coefficient of a node captures the tendency to which the neighboring nodes of a node are interconnected (Watts & Strogat, 1998), and it was computed as follows (Onnela et al., 2005)

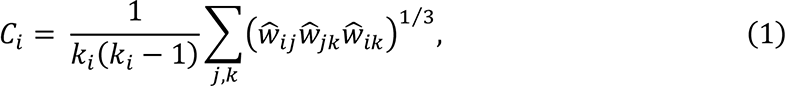

Where *Ci* corresponds to the clustering coefficient of node *i*, *ki* is the number of edges connected to node *i*, *wij* corresponds to the strength of a connection between nodes *i* and *j*, and *Ŵ*_*ij*_ represents the weight scaled by the highest weight in the network (*Ŵ*_*ij*_ =*W*_*ij*_/max (*W*)). Then, the global-level clustering coefficient was calculated by averaging the nodal-level clustering coefficient across all nodes.

The shortest path length represents the lowest cost of information moving between them. Thus, for weighted FC matrices, the nodal-level shortest path length is defined as

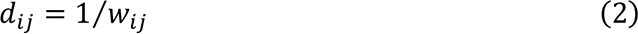

The global-level characteristic path length is given by (Percha et al., 2005).

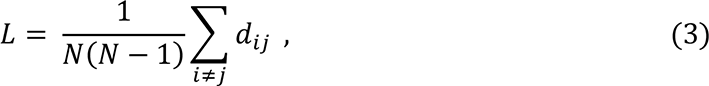

The commonly used formula to quantify small worldness was proposed by Humphries and Gurney (2008) (See SI Methods for detailed description), but it has been criticized (Bassett & Bullmore, 2017; Papo et al., 2016; Telesford et al., 2011) because it is strongly driven by network clustering and density of the graph. Recently, Telesford et al. (2011) established a new small-worldness index separately comparing network clustering to a regular network and path length to a random network. In the present study, we used an alternative index (*φ*) proposed by Muldoon et al. (2016) to investigate whether or not neonates’ brains at birth show a similar functional small-world architecture to the adult brain. That function follows Telesford and colleagues’ measure (2011) and, at the same time, takes variations in network density and connection strengths into account. The *φ* is computed according to the following equation

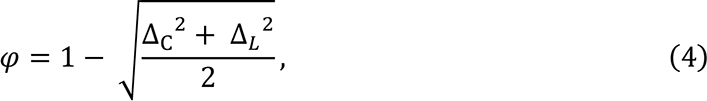

where

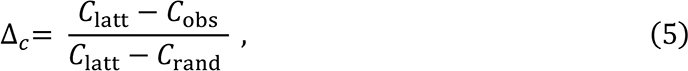

and

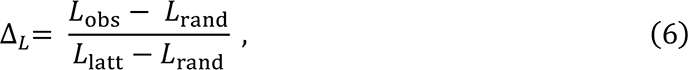

where latt/obs/rand represent lattice/observed/random networks, the *Δc* and *ΔL* represent the fractional deviation of the *Cobs* or *Lobs* from its respective lattice and random networks constructed with the same number of nodes and the same degree distribution (null model). Thus, *Δc* and *ΔL* both range from 0 to 1, and a higher value means the observed networks have a lower clustering coefficient and a higher characteristic path length. The *φ* ranges from 0 to 1, and a higher *φ* value reflects the observed networks exhibiting stronger small-world architecture.

The normalized small-world properties, *Δc*, *ΔL* and *φ*, were then averaged across all 21 thresholds for further statistical analyses. In addition, we also calculated the mean strength of FC, which can be a confound in between-group difference of small-world properties (van den Heuvel et al., 2017). The mean FC was calculated for each participant by averaging all positive edges.

#### Nodal efficiency

To quantify the communication efficiency at individual brain node, we used nodal efficiency (*Ei*), which is defined as follows (Achard & Bullmore, 2007):

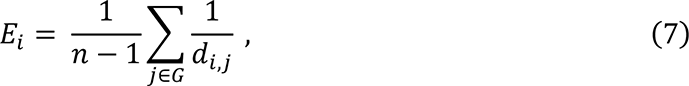

where *Ei* is the mean of the inverse shortest path length from node *i* to the other nodes. Brain node having high *Ei* exhibits a high level of efficiency in communicating with the rest of the brain.

### Statistical analyses

#### Comparison in functional brain architecture at the global level

*Adults versus neonates* To investigate how the functional brain architecture in neonates differs from that of adults, we compared small-world properties in the adults and neonate groups. General linear models (GLMs) were used to test the difference in *φ*, normalized clustering coefficient (*Δc*) or normalized characteristic path length (*ΔL*) between adults and each neonate group while controlling for mean FD and FC, which are significantly different between the adults and neonate groups (SI, Figure S1 and S2a).

*Full-term versus preterm neonates* To investigate the effect of premature birth on the development of functional architecture at the global level, we compared the normalized small-world properties (*φ*, *Δc* and *ΔL*) between the full-term neonates and preterm neonates at/before TEA. GLMs were used to detect the difference between full-term neonates and preterm neonates at or before TEA while controlling for mean FD and FC.

*Preterm neonates scanned at TEA versus preterm neonates before TEA* A smaller sample of 33 preterm neonates scanned both before and at TEA was used to investigate how small- world architecture develops in preterm neonates from before to reaching TEA. Linear mixed- effect models were applied to detect the difference in small-world properties (*φ*, Δ_*C*_ or Δ_*L*_) between preterm neonates scanned at and before TEA while controlling for mean FD and FC.

#### Comparison in nodal efficiency between full-term and preterm neonates

To investigate the effect of premature birth on efficiency of regional communication, GLMs were used to detect any differences in nodal efficiency between full-term neonates and preterm neonates at or before TEA, while controlling for mean FD and FC. Correction for the false discovery rate was applied to all statistical results at a threshold of *p* < 0.05.

### Validation analyses

The main global-level network analyses in the current study were based on weighted FC matrices and the algorithms proposed by Muldoon et al., (2016) (See SI Methods for details on definition and algorithms). To ensure that our results were not driven by specific choice of network types and algorithms, we also calculated and compared small-world architecture using 1) binary FC matrices and algorithms proposed by Muldoon et al. (2016); 2) weighted FC matrices and algorithms proposed by Humphries and Gurney (2008); and 3) binary FC matrices and algorithms proposed by Humphries and Gurney (2008). In addition, given how graph theoretical analysis results may be driven by specific parcellations, we tested the reproducibility of our results using another parcellation, the UNC-0-1-2 atlas with 223 regions (Shi et al., 2018), as a recent study (Luppi & Stamatakis, 2020) showed a parcellation with about 200 regions may be the most representative for functional network (See SI Methods for details on node definition).

## Results

### Comparison of neonate and adult functional small-world architecture

GLMs comparing the adults and full-term neonates, including head motion and mean FC as covariates, showed significant main effects of groups for *φ* (*F* (1, 450) = 74.69, *p* < 0.0001) and *Δc* (*F* (1, 450) = 167.78, *p* < 0.0001), which was driven by significantly higher *φ* and lower *Δc* in the adults relative to full-term neonates, respectively (Figure 3b). We observed similar results when comparing the adults and preterm neonate groups. For preterm neonates scanned at TEA, we found significant main effects of group for *φ* (*F* (1, 244) = 184.98, *p* < 0.0001) and *Δc* (*F* (1, 244) = 141.69, *p* < 0.0001). The adults had significantly higher *φ* and lower *Δc* relative to preterm neonates at TEA (Figure 3b). Similarly, for the preterm neonates before TEA, we found significant main effects of group for *φ* (*F* (1, 242) = 256.97, *p* < 0.0001), and *Δc* (*F* (1, 242) = 32.43, *p* < 0.0001), which were driven by significantly higher *φ* and lower *Δc* in the adults (Figure 3b). In addition, we observed a significant main effect for *ΔL* (*F* (1, 242) = 53.78, *p* < 0.0001), which was driven by significantly lower *ΔL* in the adults. In summary, as expected significantly higher small-world propensity/small worldness and clustering coefficient were consistently observed in the adults relative to neonate groups using different algorithms and types of FC matrices, which were controlled for differences in functional connectivity and head motion between adult and infant groups (SI, Figure S6).

**Figure 3.**
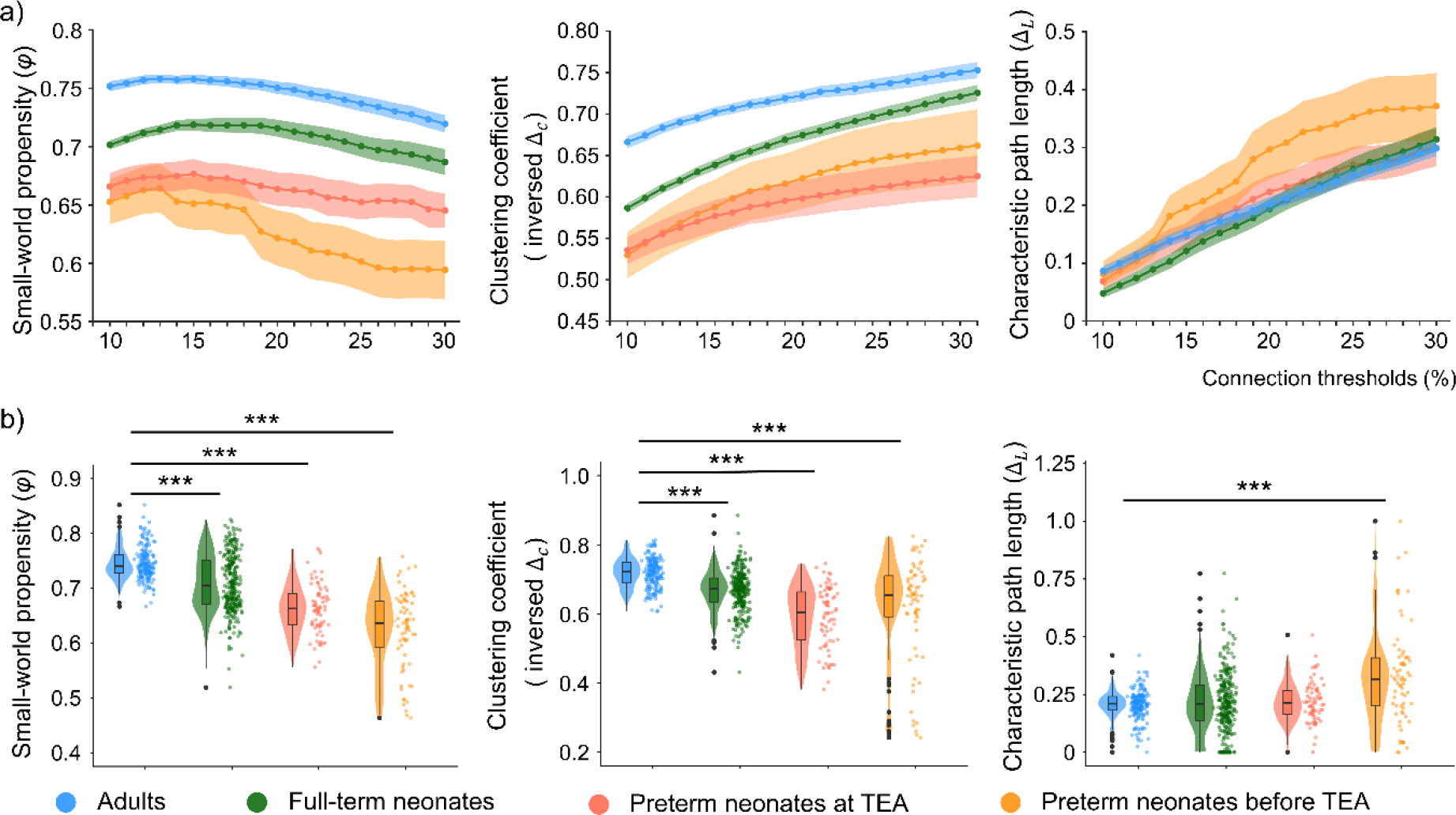
The development of small-world architecture in the neonates relative to adults. a) small-world properties at all tested thresholds (from 10% to 30% in 1% increases) and b) small-world properties in the neonate groups relative to adults. The small-world propensity, clustering coefficient, and characteristic path length shown in b) were averaged across all thresholds. Higher *φ*/inversed *ΔC*/*ΔL* represent higher small-world propensity, higher clustering coefficient and longer characteristic path length. The inversed *ΔC* was calculated by 1 - *ΔC* and for visualization purposes only and the statistical analyses were conducted using *ΔC*. The violin plots show the distribution of the data. The line that divides the box into two parts represents the median of the data. The ends of the box represent the upper (Q3) and lower (Q1) quartiles of the data. The extreme lines of the boxplot show Q3 + 1.5 * interquartile to Q1 – 1.5 * interquartile range. The black dots beyond the extreme lines show potential outliers. Abbreviations: TEA, term-equivalent age. *** = *p* < 0.0005.

### Small-world organization in neonates

When a threshold value of *φ* = 0.6 (Muldoon et al., 2016) was applied to detect whether neonates exhibited small-world architecture or not, the threshold was met or surpassed in 100% of the adults, 97.8% of full-term neonates, 90.3% of preterm neonates at TEA and 72.9% of preterm neonates before TEA (Table 2). Although capturing only a summary level view, this result suggested that small-world organization is present in a large proportion of neonates before TEA in the majority/all preterm neonates at term equivalent age or born at full term age. In the next sections, we perform more fine-grained analyses and group comparisons to unravel the effect of prematurity and early infant age on small-world organization.

**Table 2.** The percentages of participants exhibiting strong small-world architecture in the adults and neonate groups.

| | $\phi > 0.6$ | | $\sigma > 1$ | |
| --- | --- | --- | --- | --- |
|  | Weighted | Binary | Weighted | Binary |
| Adults | 100.0% | 100.0% | 92.0% | 94.3% |
| Full-term neonates | 97.8% | 97.8% | 83.1% | 90.3% |
| Preterm neonates at TEA | 90.3% | 94.4% | 76.4% | 93.1% |
| Preterm neonates before TEA | 72.9% | 71.4% | 55.7% | 74.3% |

Abbreviations: PMA, postmenstrual age in weeks; $\phi$ , small-world propensity calculated according to Muldoon et al., (2016); $\sigma$ , small-worldness calculated according to Humphries and Gurney (2008).

### The effect of premature birth on the development of small-world architecture

When the effects prematurity and early infant age where collapsed, in the comparison between full-term neonates and preterm neonates before TEA, we found significant main effects of group for *φ* (*F* (1, 344) = 86.37, *p* < 0.0001) and *ΔL* (*F* (1, 344) = 43.21, *p* < 0.0001), driven by significantly higher *φ* and lower *ΔL* in the full-term neonates (Figure 4a). These results suggested that functional architecture has higher small worldness and higher integration in full-term neonates relative to preterm neonates before TEA at birth.

**Figure 4.**
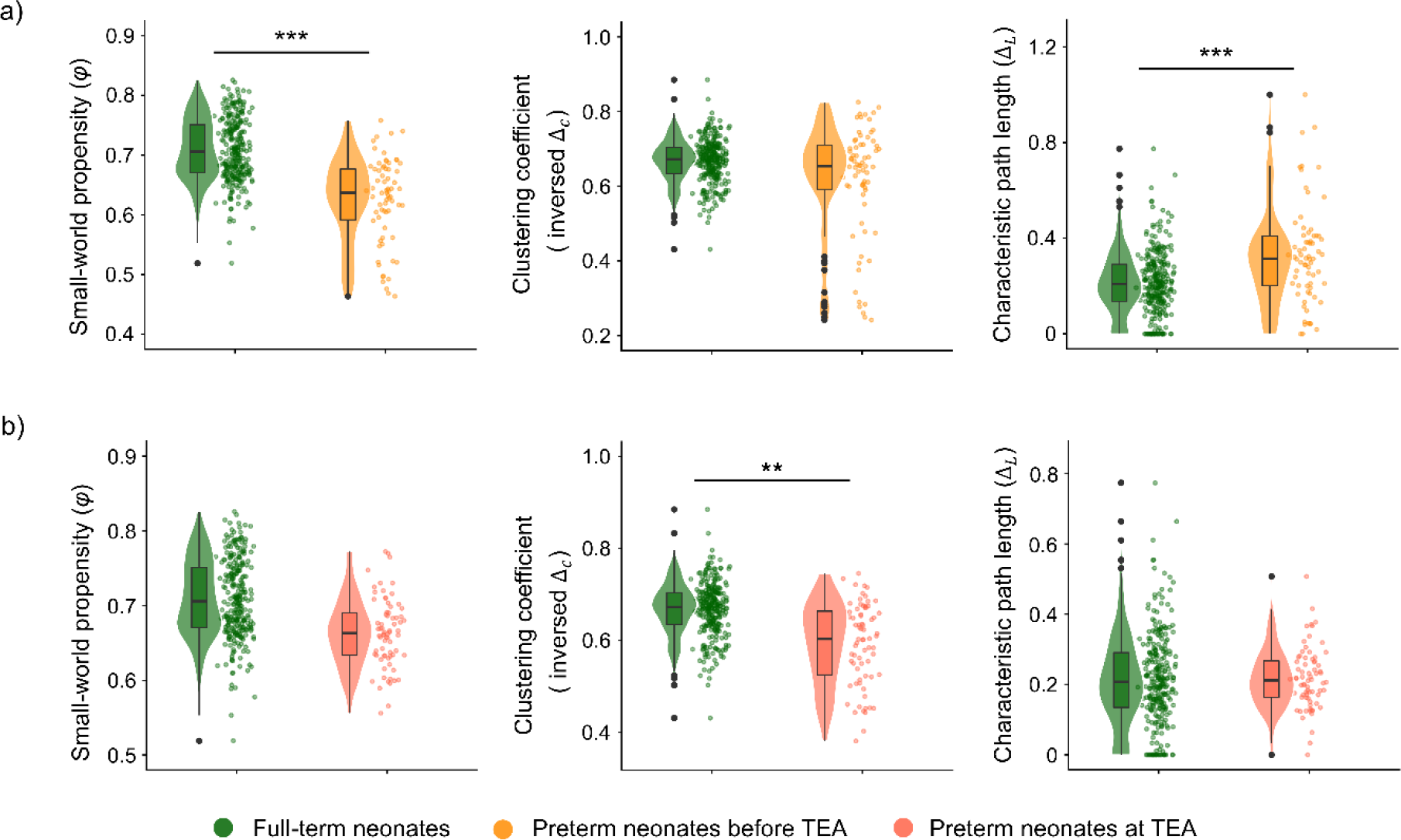
Effects of premature birth on the development of small-world architecture. a) small-world properties in the full-term neonates relative to preterm neonates before TEA; b) small-world properties in the full-term neonates relative to preterm neonates at TEA. Full-term neonates had significantly higher small-world propensity and lower normalized path length relative to preterm neonates before TEA, independent of mean functional connectivity. Full-term neonates had significantly higher clustering coefficient relative to preterm neonates at TEA. Higher *φ*/inversed *ΔC*/*ΔL* represent higher small- world propensity, higher clustering coefficient and longer characteristic path length. The inversed *ΔC* was calculated by 1 - *ΔC* and for visualization purposes only and the statistical analyses were conducted using *ΔC*. The violin plots show the distribution of the data. The line that divides the box into two parts represents the median of the data. The ends of the box represent the upper (Q3) and lower (Q1) quartiles of the data. The extreme lines of the boxplot show Q3 + 1.5 * interquartile to Q1 – 1.5 * interquartile range. The black dots beyond the extreme lines show potential outliers. Abbreviations: TEA, term-equivalent age. *** = *p* < 0.0005; ** = *p* < 0.005.

When the effect of prematurity were considered independently of early infant age, in the comparison between full-term neonates and preterm neonates at TEA, we found a trend main effect of group for *φ* (*F* (1, 346) = 5.69, *p* = 0.018), driven by higher *φ* in the full-term neonates (Figure 4b). However, this result did not survive multiple comparisons correction. In addition, we observed a significant main effect of group for *ΔC* (*F* (1, 346) = 10.78, *p* = 0.001), driven by a significantly lower *ΔC* in full-term neonates relative to preterm neonates at TEA. These results suggested that full-term born neonates had higher segregation than preterm neonates of the same age. Similar findings were observed using different algorithms and types of FC matrices (SI, Figure S6) and another parcellation (SI Results, Figure S8b).

### The effect of neonate age on the development of small-world architecture

When we investigated the effect of early infant age, independently of prematurity, by comparing the same preterm neonate group that was scanned twice - before and at TEA - we found a significant main effect of neonate age on *φ* (*F* (1, 40.21) = 13.23, *p* = 0.001), which was driven by significantly higher small-world propensity in preterm at TEA relative to the same group before TEA (Figure 5b). In addition, we found that the percentage of neonates exhibiting small-world architecture increased from 66.7% (22 out of 33) to 90.9% (30 out of 33) when they reached TEA. These results suggested that functional small-world architecture develops significantly with early infant age as premature infants reach TEA. When small-world properties are considered at a fine-grain level, we see that significant effects of premature birth remain when infants reach term equivalent age.

**Figure 5.**
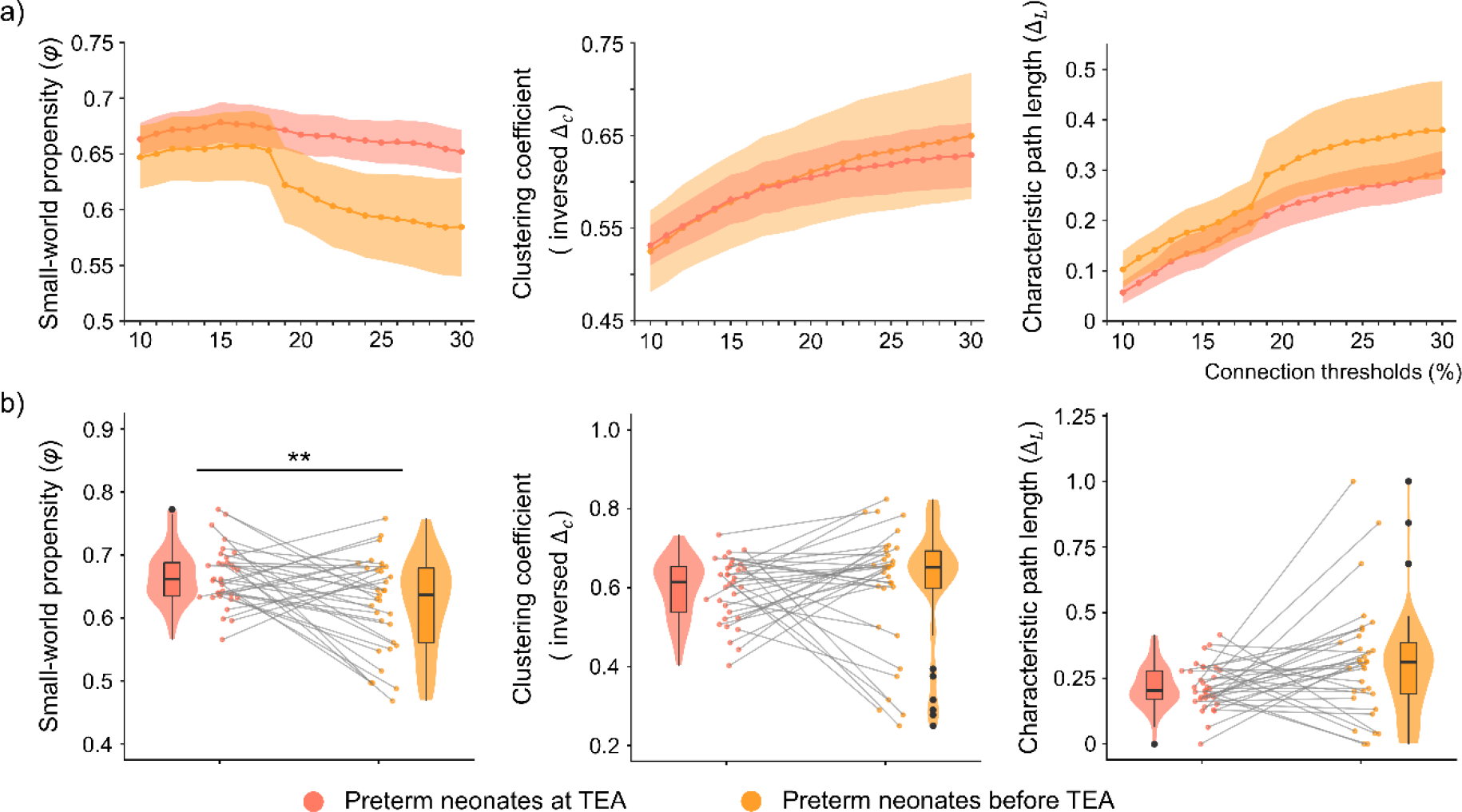
The development of small-world architecture in preterm neonates up to term- equivalent age. a) small-world properties at all tested thresholds (from 10% to 30% in 1% increases) and b) small-world properties in the preterm neonates at TEA relative to the same neonates scanned before TEA. The small-world propensity, clustering coefficient and characteristic path length shown in b) were averaged across all thresholds. Higher *φ*/inversed *ΔC*/*ΔL* represent higher small-world propensity, higher clustering coefficient and longer characteristic path length. The inversed *ΔC* was calculated by 1 - *ΔC* and for visualization purposes only and the statistical analyses were conducted using *ΔC*. The violin plots show the distribution of the data. The line that divides the box into two parts represents the median of the data. The ends of the box represent the upper (Q3) and lower (Q1) quartiles of the data. The extreme lines of the boxplot show Q3 + 1.5 * interquartile to Q1 – 1.5 * interquartile range. The black dots beyond the extreme lines show potential outliers. Abbreviations: TEA, term-equivalent age. ** = *p* < 0.005.

### The effect of premature birth and early infant age on the development of nodal efficiency

The collapsed effects of premature birth and infant age were first considered. The GLMs comparing nodal efficiency between the full-term neonates and preterm neonates before TEA, while controlling mean FC and FD, showed the full-term neonates had significantly higher nodal efficiency in 9 networks: sensorimotor mouth network (100% of nodes), followed by the sensorimotor hand network (56.67%), the dorsal attention (45.45%), the cingular opercular (42.86%), the frontoparietal control (41.67%), the salience (38.89%), the auditory (23.08%), the default mode (22.64%), and the visual network (18.52%) (Figure 6a, Table 3) (Please see SI for further supporting results, Figure S4).

**Figure 6.**
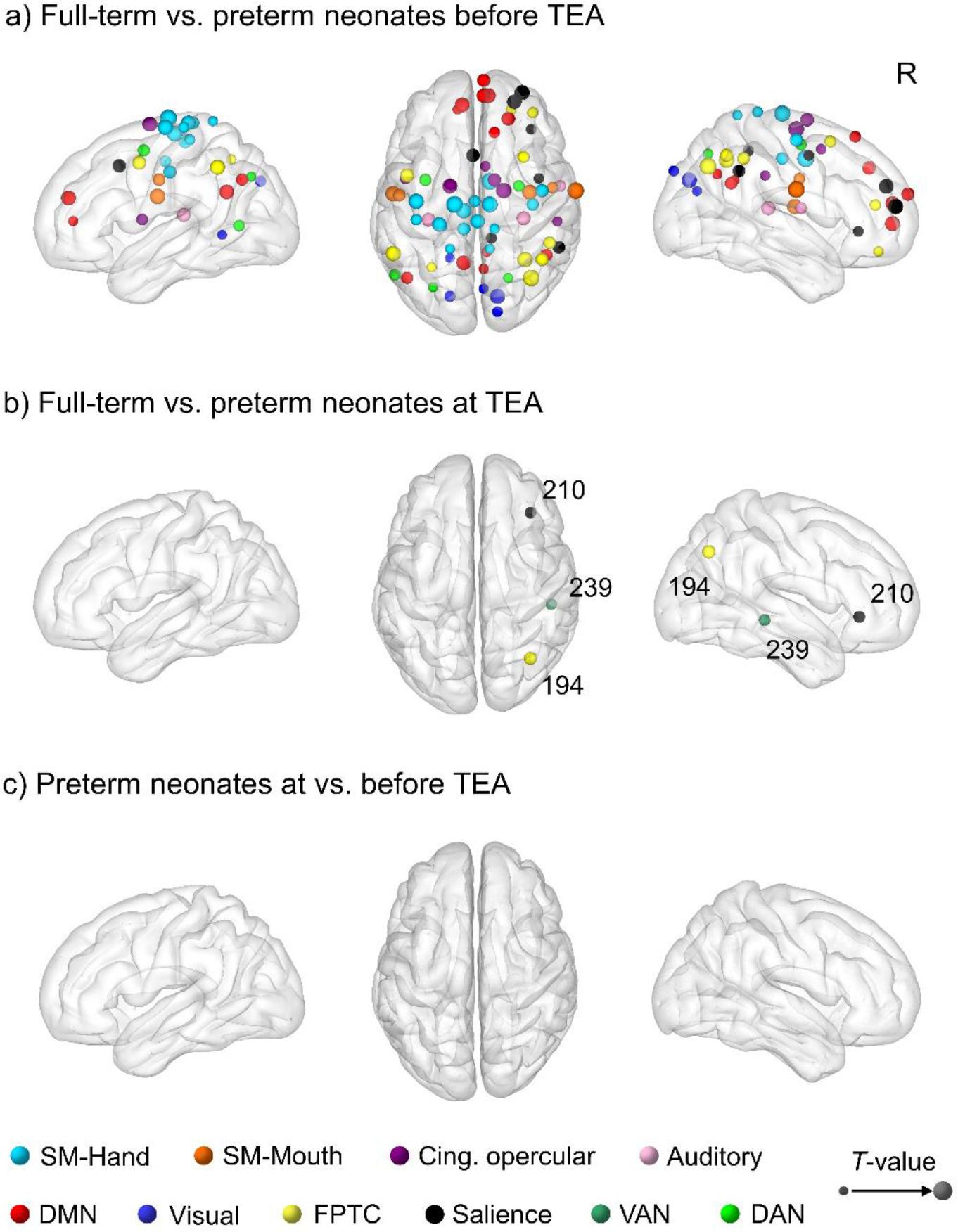
Effects of premature birth on nodal efficiency. a) nodes showing higher nodal efficiency in the full-term neonates relative to preterm neonates before TEA; b) nodes showing higher nodal efficiency in the full-term neonates relative to preterm neonates at TEA; and c) no significant difference between preterm neonates scanned at TEA and the same neonates scanned before TEA. The size of the node represents *T*-value. Abbreviations: SM-Hand, sensory/somatomotor hand network; SM-Mouth, sensory/somatomotor mouth network; Cing. opercular, cingulo-opercular task control network; DMN, Default mode network; FPTC, frontoparietal task control network; VAN, ventral attention network; DAN, dorsal attention network; R, right.

**Table 3.** The percentage of nodes showing significantly higher nodal efficiency in each network in the full-term neonates relative to preterm neonates before TEA.

| Networks | Percentage of significant nodes |
| --- | --- |
| SM-Mouth | 100.00% (5/5) |
| SM-Hand | 56.67% (17/30) |
| DAN | 45.45% (5/11) |
| Cing. opercular | 42.86% (6/14) |
| FPTC | 41.67% (10/24) |
| Salience | 38.89% (7/18) |
| Auditory | 23.08% (3/13) |
| DMN | 22.64% (12/53) |
| Visual | 18.52% (5/27) |
| Subcortical | 0.00% (0/13) |
| VAN | 0.00% (0/9) |

Abbreviations: SM-Hand, sensory/somatomotor hand network; SM-Mouth, sensory/somatomotor mouth network; Cing. opercular, cingulo-opercular task control network; DMN, Default mode network; FPTC, frontoparietal task control network; VAN, ventral attention network; DAN, dorsal attention network.

Second, we considered the effect of premature birth independently of infant age. The GLMs comparing nodal efficiency between the full-term neonates and preterm neonates at TEA showed that the full-term neonates had higher nodal efficiency in the right angular gyrus (node 194, frontoparietal control network), the orbital part of right inferior frontal gyrus (node 210, salience network) and the right middle temporal gyrus (node 239, ventral attention network) (Figure 6b). Significant differences in these nodes were also observed relative to preterm neonates before TEA. Third, when we considered the effect infant age independently of prematurity by comparing nodal efficiency between the same preterm neonates scanned before and at TEA no significant main effects were found (Figure 6c).

## Discussion

We investigated the development of brain’s functional small-world architecture in neonates and how that architecture may be altered by premature birth. To the best of our knowledge, this study is the first fMRI study to investigate the effect of prematurity before term equivalent age both on the global small-world properties and long-range regional communication. The key novel finding of this study is that premature neonates before term equivalent age show significant differences in small-world brain architecture and in the efficiency of regional communication relative to full term born neonates. These differences are driven by both sensory and higher-order networks, with the somatomotor, the dorsal attention, the cingulo-opercular and the frontoparietal control networks carrying the largest effects.

### Comparison of neonate and adult functional architectures

As expected, adults had stronger small-worldness values, including higher clustering coefficient than all neonate groups, suggesting more efficient and segregated functional brain architecture relative to neonates. It is worth noting that differences in functional architecture between adults and neonate groups and across neonate groups were independent of any differences in functional connectivity or head motion. These findings are consistent with previous studies showing that the functional organization of the brain develops rapidly from birth onwards, with functional brain networks gradually becoming more cohesive (Cabral et al., 2022; Sherman et al., 2014; Gao et al., 2009) and specialized (Wen et al., 2019; Scheinost et al., 2016; Fransson et al., 2011) after birth.

### Effect of premature birth

At birth, preterm neonates were less likely to exhibit functional small-world architecture relative to full-term neonates. However, while a small-world architecture was not observed in some preterm infants before TEA (27%) and at TEA (< 10%), it was interesting to see that from a global view the majority of the preterm infants both before and at TEA exhibited small-world organization, even though its properties were altered relative to full-term neonates.

A more refined picture emerged when group differences of small-world properties across infant groups were considered. In infants born before TEA, premature birth was associated with significantly weaker small-world organization and higher path length, or lower neural integration, relative to full-term birth. Our finding differs from two previous studies, which reported stronger small-world properties in preterm neonates before TEA (Cao et al., 2017; Omidvarnia et al., 2014), a discrepancy likely due the small sample sizes (N = 11 to 23) in these previous studies. Our results are consistent with previous studies showing that the strength of functional connectivity increases with gestational age (Canini et al., 2020; Keunen et al., 2017; Van Den Heuvel et al., 2015; Thomason et al., 2015, 2013), providing further evidence for the importance of the third trimester for infant brain development.

As preterm neonates reached TEA, we found that functional organization matured significantly, including significantly stronger small wordless and shorter path length (higher neural integration), as observed in a smaller group of infants (N = 33) that were scanned longitudinally at both preterm and term-equivalent age time points. Nevertheless, some disruptions of small-world organization persisted in premature neonates at TEA relative to full-term born counterparts. We found a trend on weaker small world organization and significantly smaller clustering coefficient (lower neural segregation), in preterms at TEA. These results are consonant with previous studies showing that prematurity was associated with lower within-network functional connectivity and less developed functional networks in preterms at TEA relative to full-term neoneates (Hu et al., 2022; Eyre et al., 2021; Bouyssi-Kobar et al., 2019; Gozdas et al., 2018; Smyser et al., 2010). Premature birth has also been associated with decreased segregation, or lower network modularity observed in preterms at TEA relative to full-term born neonates (Scheinost et al., 2016).

To shed light on brain regions and networks that drive these global functional architecture differences across infant groups, we assessed nodal efficiency across 217 nodes comprising 11 brain networks with comprehensive brain coverage. Premature birth before TEA, was associated with decreased nodal efficiency in 32% of nodes, distributed across 9/11 brain networks. Relative to full-term neonates, premature neonates born before TEA showed significantly lower nodal efficiency in sensory-driven and higher-order networks with the somatomotor, dorsal attention, cingulo-opercular, and frontoparietal control network showing significantly reduced efficiency in over 40% of nodes. The somatomotor network encompassing regions involved in mouth and hand movement showed significantly reduced efficiency in 100% and 57% of the nodes respectively. Foetuses can receive somatomotor input from their muscles and limb movements in utero, which is key for early development of movement abilities. Our data suggests that the somatomotor networks undergoe the largest development of communication efficiency in the 3rd trimester relative to all other brain networks.

By contrast, the subcortical and ventral attention networks showed no differences between preterms before TEA and full-term neonates, suggesting that efficiency of regional communication in these two networks does not change significantly in the third trimester. Consistent with this, the visual network showed the efficiency in the fewest nodes (< 20%) associated with prematurity before TEA. As all preterm infants born before TEA and full-term born infants were scanned at or very shortly after birth, these neonates would have had very limited extra-uterine sensory experience. Our findings are consistent with the limited visuo-sensory experience of neonates and suggest that visually-driven brain networks develop at a lesser pace than others in the 3^rd^ trimester, even in infants who are exposed to extra- uterine environments before TEA, and rather pick up developmental page substantially post birth, once infants receive continuous visual input (Dubois et al., 2021; Ko et al., 2013; Colonnese et al., 2010). This explanation is supported by the finding that the number of connections between visual regions and the rest of the brain increased with neonate age in preterm neonates, and at term-equivalent age preterm neonates had more connections in visual regions than full-term neonates (Fenn-Moltu et al., 2023).

By contrast to premature neonates before TEA, premature neonates at TEA showed significantly reduced efficiency in only three nodes (i.e., 32% vs 1.4%) belonging to the salience, visual attention network and frontoparietal control networks. In addition to belonging to networks that support attention and executive function, these nodes fall within the right angular, inferior frontal gyrus and middle temporal gyrus, regions that are related to language processing (Friederici, 2011; Perani et al., 2011; Hickok & Poeppel, 2007).

Therefore, disruption of nodal efficiency in these regions associated with prematurity in term equivalent neonates may underlie the significant risks for language development delays that are associated with premature birth. Children born prematurely are at risk for problems in language acquisition, word comprehension, and the development of receptive language skills (Vandormael et al., 2019; Rushe et al., 2010; Guarini et al., 2009).

### Methodological considerations

Although some previous studies have reported a negative association of prematurity with small-world properties (Cao et al., 2017), others have not (Bouyssi-Kobar et al. 2019; Omidvarnia et al., 2014). This inconsistency is likely due to methodological differences in the type of data acquired, analyses method, and small sample sizes in these previous studies.

Here we were enabled by the high quality dHCP dataset to employ a uniquely large sample (N = 278) of high temporal and spatial resolution infant rs-fMRI data, and accurate week-by-week structural templates of the developing infant brain, which ensured higher sensitivity to detect brain functional architecture changes in early infancy. Given these methodological strengths and clear results, we believe this study helps to resolve previous inconsistent findings on the development of functional brain architecture in neonates. We note that the preterm group is not well representative of extreme prematurity (<28 weeks) / very low birth weight (<1500g), because the mean GA of the preterm group was 32 weeks. Extreme prematurity could be associated with different functional architectures at term-equivalent age, as well as during the preterm period, compared with infants with higher GAs, and will be investigated in future studies.

### Summary

In summary, our findings show that premature neonates before term equivalent age show significant differences in small-world brain architecture and efficiency of regional communication relative to full term born neonates. These differences are driven by both sensory and higher-order networks, and show an effect hierarchy, with the somatomotor networks carrying the largest spatial effect and the visual network the smallest. These findings emphasize the importance of the third trimester in developing the functional architecture of the brain architecture. Our results demonstrate that small-world organization and the efficiency of regional communication increases gradually with perinatal age in the 3^rd^ trimester up to full-term birth, rather than undergoing drastic maturation steps in this period. Furthermore, they suggest that, for the most part, small-world brain architecture and regional communication efficiency develops in healthy-born premature neonates by term-equivalent age, according to a pre-programmed developmental trajectory despite of prematurity.

Nevertheless, in preterm born neonates at term equivalent age prematurity was associated with disrupted small-world architecture and reduced efficiency of regional communication in networks related with high-order cognition, including language. Adding to a growing literature (Fenn-Moltu et al., 2023; Vanes et al., 2023; Hu et al., 2022; Della et al., 2021; Eyre et al., 2021; Bouyssi-Kobar et al., 2019; Scheinost et al., 2016), our results shed light on disrupted brain mechanisms that may underlie the significant risks for neurodevelopmental and psychiatric problems in later life (Nosarti et al., 2012; Saigal et al., 2008; Marlow et al., 2005; Bhutta et al., 2002), that are associated with premature birth. These findings improve understanding of the ontogeny of functional small-world architecture and efficiency of neural communication across specific brain networks.

## Acknowledgments

Neonate data were provided by the developing Human Connectome Project (http://www.developingconnectome.org/), KCL-Imperial-Oxford Consortium funded by the European Research Council under the European Union Seventh Framework Programme (FP/2007-2013) / ERC Grant Agreement no. [319456]. We are grateful to the families who generously supported this trial. Adult data were provided [in part] by the Human Connectome Project (http://www.humanconnectomeproject.org), WU-Minn Consortium (Principal Investigators: David Van Essen and Kamil Ugurbil; 1U54MH091657) funded by the 16 NIH Institutes and Centers that support the NIH Blueprint for Neuroscience Research; and by the McDonnell Center for Systems Neuroscience at Washington University.

## References

1. Achard, S., & Bullmore, E. (2007). Efficiency and cost of economical brain functional networks. PloS computational biology, 3(2), e17.

2. Allievi, A. G., Arichi, T., Tusor, N., Kimpton, J., Arulkumaran, S., Counsell, S. J., … & Burdet, E. (2016). Maturation of sensori-motor functional responses in the preterm brain. Cerebral Cortex, 26(1), 402–413.

3. Asis-Cruz, D., Bouyssi-Kobar, M., Evangelou, I., Vezina, G., & Limperopoulos, C. (2015). Functional properties of resting state networks in healthy full-term newborns. Scientific reports, 5(1), 1–15.

4. Barttfeld, P., Uhrig, L., Sitt, J. D., Sigman, M., Jarraya, B., & Dehaene, S. (2015). Signature of consciousness in the dynamics of resting-state brain activity. Proceedings of the National Academy of Sciences, 112(3), 887–892.

5. Bassett, D. S., & Bullmore, E. D. (2006). Small-world brain networks. The neuroscientist, 12(6), 512–523.

6. Bassett, D. S., & Bullmore, E. T. (2017). Small-world brain networks revisited. The Neuroscientist, 23(5), 499–516.

7. Bhutta, A. T., Cleves, M. A., Casey, P. H., Cradock, M. M., & Anand, K. J. (2002). Cognitive and behavioral outcomes of school-aged children who were born preterm: a meta-analysis. Jama, 288(6), 728–737.

8. Bouyssi-Kobar, M., De Asis-Cruz, J., Murnick, J., Chang, T., & Limperopoulos, C. (2019). Altered functional brain network integration, segregation, and modularity in infants born very preterm at term-equivalent age. The Journal of pediatrics, 213, 13–21.

9. Bullmore, E., & Sporns, O. (2009). Complex brain networks: graph theoretical analysis of structural and functional systems. Nature reviews neuroscience, 10(3), 186–198.

10. Bullmore, E., & Sporns, O. (2012). The economy of brain network organization. Nature reviews neuroscience, 13(5), 336–349.

11. Cabral, L., Zubiaurre-Elorza, L., Wild, C. J., Linke, A., & Cusack, R. (2022). Anatomical correlates of category-selective visual regions have distinctive signatures of connectivity in neonates. Developmental Cognitive Neuroscience, 58, 101179.

12. Canini, M., Cavoretto, P., Scifo, P., Pozzoni, M., Petrini, A., Iadanza, A., … & Della Rosa, P. A. (2020). Subcortico-cortical functional connectivity in the fetal brain: a cognitive development blueprint. Cerebral cortex communications, 1(1), tgaa008.

13. Cao, M., He, Y., Dai, Z., Liao, X., Jeon, T., Ouyang, M., … & Huang, H. (2017). Early development of functional network segregation revealed by connectomic analysis of the preterm human brain. Cerebral cortex, 27(3), 1949–1963.

14. Colonnese, M. T., Kaminska, A., Minlebaev, M., Milh, M., Bloem, B., Lescure, S., … & Khazipov, R. (2010). A conserved switch in sensory processing prepares developing neocortex for vision. Neuron, 67(3), 480–498.

15. Cooney, J. W., & Gazzaniga, M. S. (2003). Neurological disorders and the structure of human consciousness. Trends in Cognitive Sciences, 7(4), 161–165.

16. Dall’Orso, S., Arichi, T., Fitzgibbon, S. P., Edwards, A. D., Burdet, E., & Muceli, S. (2022). Development of functional organization within the sensorimotor network across the perinatal period. Human brain mapping, 43(7), 2249–2261.

17. Deco, G., Tononi, G., Boly, M., & Kringelbach, M. L. (2015). Rethinking segregation and integration: contributions of whole-brain modelling. Nature Reviews Neuroscience, 16(7), 430–439.

18. Doria, V., Beckmann, C. F., Arichi, T., Merchant, N., Groppo, M., Turkheimer, F. E., … & Edwards, A. D. (2010). Emergence of resting state networks in the preterm human brain. Proceedings of the National Academy of Sciences, 107(46), 20015–20020.

19. Dubois, J., Alison, M., Counsell, S. J., Hertz-Pannier, L., Hüppi, P. S., & Benders, M. J. (2021). MRI of the neonatal brain: a review of methodological challenges and neuroscientific advances. Journal of Magnetic Resonance Imaging, 53(5), 1318–1343.

20. Duclos, C., Nadin, D., Mahdid, Y., Tarnal, V., Picton, P., Vanini, G., … & Blain-Moraes, S. (2021). Brain network motifs are markers of loss and recovery of consciousness. Scientific reports, 11(1), 1–13.

21. Edelman, G. M. (1989). The remembered present: a biological theory of consciousness. Basic Books.

22. Eyre, M., Fitzgibbon, S. P., Ciarrusta, J., Cordero-Grande, L., Price, A. N., Poppe, T., … & Edwards, A. D. (2021). The Developing Human Connectome Project: typical and disrupted perinatal functional connectivity. Brain, 144(7), 2199–2213.

23. Fenn-Moltu, S., Fitzgibbon, S. P., Ciarrusta, J., Eyre, M., Cordero-Grande, L., Chew, A., … & Batalle, D. (2023). Development of neonatal brain functional centrality and alterations associated with preterm birth. Cerebral Cortex, 33(9), 5585–5596.

24. Fitzgibbon, S. P., Harrison, S. J., Jenkinson, M., Baxter, L., Robinson, E. C., Bastiani, M., … & Andersson, J. (2020). The developing Human Connectome Project (dHCP) automated resting-state functional processing framework for newborn infants. Neuroimage, 223, 117303.

25. Fransson, P., Åden, U., Blennow, M., & Lagercrantz, H. (2011). The functional architecture of the infant brain as revealed by resting-state fMRI. Cerebral cortex, 21(1), 145–154.

26. Friederici, A. D. (2011). The brain basis of language processing: from structure to function. Physiological reviews, 91(4), 1357–1392.

27. Gao W, Zhu HT, Giovanello KS, et al. Evidence on the emergence of the brain’s default network from 2-week-old to 2-year-old healthy pediatric subjects. Proceedings of the National Academy of Sciences . 2009;106(16):6790–6795.

28. Gao, W., Alcauter, S., Elton, A., Hernandez-Castillo, C. R., Smith, J. K., Ramirez, J., & Lin, W. (2015). Functional network development during the first year: relative sequence and socioeconomic correlations. Cerebral cortex, 25(9), 2919–2928.

29. Glasser, M. F., Sotiropoulos, S. N., Wilson, J. A., Coalson, T. S., Fischl, B., Andersson, J. L., … & Wu-Minn HCP Consortium. (2013). The minimal preprocessing pipelines for the Human Connectome Project. Neuroimage, 80, 105–124.

30. Godwin, D., Barry, R. L., & Marois, R. (2015). Breakdown of the brain’s functional network modularity with awareness. Proceedings of the National Academy of Sciences, 112(12), 3799–3804.

31. Gozdas, E., Parikh, N. A., Merhar, S. L., Tkach, J. A., He, L., & Holland, S. K. (2018). Altered functional network connectivity in preterm infants: antecedents of cognitive and motor impairments?. Brain Structure and Function, 223(8), 3665–3680.

32. Guarini, A., Sansavini, A., Fabbri, C., Alessandroni, R., Faldella, G., & Karmiloff-Smith, A. (2009). Reconsidering the impact of preterm birth on language outcome. Early human development, 85(10), 639–645.

33. Hallquist, M. N., & Hillary, F. G. (2018). Graph theory approaches to functional network organisation in brain disorders: A critique for a brave new small-world. Network Neuroscience, 3(1), 1–26. 3-395.

34. Hickok, G., & Poeppel, D. (2007). The cortical organization of speech processing. Nature reviews neuroscience, 8(5), 393–402.

35. Hu, H., Cusack, R., & Naci, L. (2022). Typical and disrupted brain circuitry for conscious awareness in full-term and preterm infants. Brain communications, 4(2), fcac071.

36. Humphries, M. D., & Gurney, K. (2008). Network ‘small-world-ness’: a quantitative method for determining canonical network equivalence. PloS one, 3(4), e0002051.

37. Ito, T., Brincat, S. L., Siegel, M., Mill, R. D., He, B. J., Miller, E. K., … & Cole, M. W. (2020). Task-evoked activity quenches neural correlations and variability across cortical areas. PLoS computational biology, 16(8), e1007983.

38. Keunen, K., Counsell, S. J., & Benders, M. J. (2017). The emergence of functional architecture during early brain development. Neuroimage, 160, 2–14.

39. Ko, H., Cossell, L., Baragli, C., Antolik, J., Clopath, C., Hofer, S. B., & Mrsic-Flogel, T. D. (2013). The emergence of functional microcircuits in visual cortex. Nature, 496(7443), 96–100.

40. Lemaître, H., Augé, P., Saitovitch, A., Vinçon-Leite, A., Tacchella, J. M., Fillon, L., … & Zilbovicius, M. (2021). Rest functional brain maturation during the first year of life. Cerebral Cortex, 31(3), 1776–1785.

41. Luppi, A. I., & Stamatakis, E. A. (2021). Combining network topology and information theory to construct representative brain networks. Network Neuroscience, 5(1), 96–124.

42. Luppi, A. I., Craig, M. M., Pappas, I., Finoia, P., Williams, G. B., Allanson, J., … & Stamatakis, E. A. (2019). Consciousness-specific dynamic interactions of brain integration and functional diversity. Nature communications, 10(1), 1–12.

43. Luppi, A. I., Golkowski, D., Ranft, A., Ilg, R., Jordan, D., Menon, D. K., & Stamatakis, E. A. (2021). Brain network integration dynamics are associated with loss and recovery of consciousness induced by sevoflurane. Human Brain Mapping, 42(9), 2802–2822.

44. Marlow, N., Wolke, D., Bracewell, M. A., & Samara, M. (2005). Neurologic and developmental disability at six years of age after extremely preterm birth. New England journal of medicine, 352(1), 9–19.

45. Monti, M. M., Lutkenhoff, E. S., Rubinov, M., Boveroux, P., Vanhaudenhuyse, A., Gosseries, O., … & Laureys, S. (2013). Dynamic change of global and local information processing in propofol-induced loss and recovery of consciousness. PLoS computational biology, 9(10), e1003271.

46. Muldoon, S. F., Bridgeford, E. W., & Bassett, D. S. (2016). Small-world propensity and weighted brain networks. Scientific reports, 6(1), 1–13.

47. Nosarti, C., Reichenberg, A., Murray, R. M., Cnattingius, S., Lambe, M. P., Yin, L., … & Hultman, C. M. (2012). Preterm birth and psychiatric disorders in young adult life. Archives of general psychiatry, 69(6), 610–617.

48. Omidvarnia, A., Fransson, P., Metsäranta, M., & Vanhatalo, S. (2014). Functional bimodality in the brain networks of preterm and term human newborns. Cerebral cortex, 24(10), 2657–2668.

49. Onnela, J. P., Saramäki, J., Kertész, J., & Kaski, K. (2005). Intensity and coherence of motifs in weighted complex networks. Physical Review E, 71(6), 065103.

50. Papo, D., Zanin, M., Martínez, J. H., & Buldú, J. M. (2016). Beware of the small-world neuroscientist!. Frontiers in human neuroscience, 10, 96.

51. Perani, D., Saccuman, M. C., Scifo, P., Anwander, A., Spada, D., Baldoli, C., … & Friederici, A. D. (2011). Neural language networks at birth. Proceedings of the National Academy of Sciences, 108(38), 16056–16061.

52. Percha, B., Dzakpasu, R., Żochowski, M., & Parent, J. (2005). Transition from local to global phase synchrony in small world neural network and its possible implications for epilepsy. Physical Review E, 72(3), 031909.

53. Power, J. D., Cohen, A. L., Nelson, S. M., Wig, G. S., Barnes, K. A., Church, J. A., … & Petersen, S. E. (2011). Functional network organization of the human brain. Neuron, 72(4), 665–678.

54. Rubinov, M., & Sporns, O. (2010). Complex network measures of brain connectivity: uses and interpretations. Neuroimage, 52(3), 1059–1069.

55. Rushe, T. M. (2010). Language function after preterm birth. Neurodevelopmental outcomes of preterm birth, 176-184.

56. Saigal, S., & Doyle, L. W. (2008). An overview of mortality and sequelae of preterm birth from infancy to adulthood. The Lancet, 371(9608), 261–269.

57. Salvador R, Martinez A, Pomarol-Clotet E, et al. A simple view of the brain through a frequency-specific functional connectivity measure. Neuroimage. 2008;39(1):279–289.

58. Scheinost, D., Kwon, S. H., Shen, X., Lacadie, C., Schneider, K. C., Dai, F., … & Constable, R. T. (2016). Preterm birth alters neonatal, functional rich club organization. Brain Structure and Function, 221(6), 3211–3222.

59. Schröter, M. S., Spoormaker, V. I., Schorer, A., Wohlschläger, A., Czisch, M., Kochs, E. F., … & Ilg, R. (2012). Spatiotemporal reconfiguration of large-scale brain functional networks during propofol-induced loss of consciousness. Journal of Neuroscience, 32(37), 12832–12840.

60. Schuh, A., Makropoulos, A., Robinson, E. C., Cordero-Grande, L., Hughes, E., Hutter, J., … & Rueckert, D. (2018). Unbiased construction of a temporally consistent morphological atlas of neonatal brain development. BioRxiv, 251512.

61. Seth, A. (2009). Explanatory correlates of consciousness: theoretical and computational challenges. Cognitive Computation, 1(1), 50–63.

62. Seth, A. K., Izhikevich, E., Reeke, G. N., & Edelman, G. M. (2006). Theories and measures of consciousness: an extended framework. Proceedings of the National Academy of Sciences, 103(28), 10799–10804.

63. Sherman, L. E., Rudie, J. D., Pfeifer, J. H., Masten, C. L., McNealy, K., & Dapretto, M. (2014). Development of the default mode and central executive networks across early adolescence: a longitudinal study. Developmental cognitive neuroscience, 10, 148–159.

64. Shi, F., Salzwedel, A. P., Lin, W., Gilmore, J. H., & Gao, W. (2018). Functional brain parcellations of the infant brain and the associated developmental trends. Cerebral Cortex, 28(4), 1358–1368.

65. Smyser, C. D., Inder, T. E., Shimony, J. S., Hill, J. E., Degnan, A. J., Snyder, A. Z., & Neil, J. J. (2010). Longitudinal analysis of neural network development in preterm infants. Cerebral cortex, 20(12), 2852–2862.

66. Telesford, Q. K., Joyce, K. E., Hayasaka, S., Burdette, J. H., & Laurienti, P. J. (2011). The ubiquity of small-world networks. Brain connectivity, 1(5), 367–375.

67. Thomason, M. E., Dassanayake, M. T., Shen, S., Katkuri, Y., Alexis, M., Anderson, A. L., … & Romero, R. (2013). Cross-hemispheric functional connectivity in the human fetal brain. Science translational medicine, 5(173), 173ra24-173ra24.

68. Thomason, M. E., Grove, L. E., Lozon Jr, T. A., Vila, A. M., Ye, Y., Nye, M. J., … & Romero, R. (2015). Age-related increases in long-range connectivity in fetal functional neural connectivity networks in utero. Developmental cognitive neuroscience, 11, 96–104.

69. Tononi, G., & Edelman, G. M. (1998). Consciousness and complexity. Science, 282(5395), 1846–1851.

70. Truzzi, A., & Cusack, R. (2023). The development of intrinsic timescales: A comparison between the neonate and adult brain. NeuroImage, 120155.

71. Uehara, T., Yamasaki, T., Okamoto, T., Koike, T., Kan, S., Miyauchi, S., … & Tobimatsu, S. (2014). Efficiency of a “small-world” brain network depends on consciousness level: a resting-state fMRI study. Cerebral cortex, 24(6), 1529–1539.

72. Van den Heuvel, M. P., de Lange, S. C., Zalesky, A., Seguin, C., Yeo, B. T., & Schmidt, R. (2017). Proportional thresholding in resting-state fMRI functional connectivity networks and consequences for patient-control connectome studies: Issues and recommendations. Neuroimage, 152, 437–449.

73. Van Den Heuvel, M. P., Kersbergen, K. J., De Reus, M. A., Keunen, K., Kahn, R. S., Groenendaal, F., … & Benders, M. J. (2015). The neonatal connectome during preterm brain development. Cerebral cortex, 25(9), 3000–3013.

74. Van Essen, D. C., Smith, S. M., Barch, D. M., Behrens, T. E., Yacoub, E., Ugurbil, K., & Wu-Minn HCP Consortium. (2013). The WU-Minn human connectome project: an overview. Neuroimage, 80, 62–79.

75. Vandormael, C., Schoenhals, L., Hüppi, P. S., Filippa, M., & Borradori Tolsa, C. (2019). Language in preterm born children: Atypical development and effects of early interventions on neuroplasticity. Neural Plasticity, 2019.

76. Vanes, L., Fenn-Moltu, S., Hadaya, L., Fitzgibbon, S., Cordero-Grande, L., Price, A., … & Nosarti, C. (2023). Longitudinal neonatal brain development and socio-demographic correlates of infant outcomes following preterm birth. Developmental Cognitive Neuroscience, 61, 101250.

77. Watts, D. J., & Strogatz, S. H. (1998). Collective dynamics of ‘small-world’networks. Nature, 393(6684), 440–442.

78. Wen, X., Zhang, H., Li, G., Liu, M., Yin, W., Lin, W., … & Shen, D. (2019). First-year development of modules and hubs in infant brain functional networks. Neuroimage, 185, 222–235.

79. Zhang, H., Shen, D., & Lin, W. (2019). Resting-state functional MRI studies on infant brains: a decade of gap-filling efforts. Neuroimage, 185, 664–684.

80. Zuo, X. N., Di Martino, A., Kelly, C., Shehzad, Z. E., Gee, D. G., Klein, D. F., … & Milham, M. P. (2010). The oscillating brain: complex and reliable. Neuroimage, 49(2), 1432–1445.

