## Supplementary material for "Typical and disrupted small-world architecture and regional communication in full-term and preterm infants": SI

**Title**

### Supplementary Methods

#### Data pre-processing

*dHCP*. The dHCP resting-state functional MRI (rs-fMRI) data were pre-processed by dHCP group using the project's in-house pipeline optimized for neonatal imaging and specifically developed for this dataset, detailed in Fitzgibbon et al. (2020). This pipeline includes: 1) motion and distortion correction: corrects for intra-volume movement artefacts and for artefacts associated with susceptibility-induced off-resonance field changes (susceptibility-by-movement artefacts) and estimates motion nuisance regressors; 2) Registration: aligns all functional images with the native T2 space and the neonatal template space, which refers to the week-specific 40-week template from the dHCP volumetric atlas (Schuh et al., 2018); 3) Temporal high-pass filter: 150s high-pass cut-off; 4) Denoising: Estimates artefact nuisance regressors and regresses all nuisance regressors from the functional data obtained from the first step.

*HCP*. The rs-fMRI data of HCP were pre-processed by HCP group using the following pipeline: 1) Distortion correction: correction of gradient-nonlinearity-induced distortion and phase-encoding-direction-induced distortion; 2) Motion correction: realigns the timeseries to correct for subject motion by using a 6 DOF FLIRT (Oxford Centre for Functional MRI of the Brain Fs Linear Registration Tool) registration of each frame to the single-band reference image; 3) Aligns the original rs-fMRI data to Montreal Neurological Institute (MNI) template space: EPI to T1w from FLIRT BBR, fine tuning of EPI to T1w with BBR-register, nonlinear T1w to MNI template; 4) Intensity normalization to mean of 10000 and bias field removal; 5) Temporal high-pass filter: 150s high-pass cut-off; 6) Denoising: removes artefactual or "bad" components using ICA-FIX to automatically. Detailed pre-processing procedure can be found in Glasser et al. (2013).

### Node definition

*Power atlas* We first aligned 226 regions of interest (ROIs) (8-mm radius spheres) in MNI space from Power et al., (2011) with 40-week dHCP T1w template. This involved: 1) trimming the dura from the 40-week dHCP T1w template, and the cerebellum from both the 40-week dHCP T1w template and MNI T1w template; 2) aligning the 40-week dHCP T1w template to MNI T1w template using non-linear registration (ANTs SyN); 3) applying the warp file generated in the last step to the ROIs in MNI space with 40-week dHCP T1w template as a reference. In the next step, we needed to align these ROIs in 40-week dHCP T1w template space with neonate native space. We inverted the func-to-template warp provided by dHCP group and applied this inverted warp to ROIs in the 40-week dHCP T1w template space. Thus, we obtained ROIs in each neonate native functional space. For adults, as the denoised HCP data had been aligned to MNI space, we used these ROIs in MNI space directly.

*UNC infant 0-1-2 atlas* The UNC infant 0-1-2 atlas was generated based on 95 infants scanned at 5 weeks, 1 year and 2 years after birth. The first release of UNC atlas (Shi et al., 2011) was generated by propagating the automated anatomical labelling (AAL) brain parcellations from adult to infant spaces based on structural similarity. To address the potential cancellation effects by averaging across heterogeneous BOLD signals within a given ROI when applying structural brain atlas in rs-fMRI studies, Shi et al., (2017) derived the first set of brain atlases by parcellating the infant brain into functionally homogeneous regions based on a novel hybrid iterative normalized cut approach. In the current study, we used the neonatal atlas from the recently released UNC infant atlas, which has 223 functionally homogenous regions.

We first aligned 223 regions of interest in MNI space provided by Shi et al., (2017) with 40-week dHCP T1w template. This involved: 1) aligning the 40-week dHCP T1w template with no dura and cerebellum to neonatal T1w image (with no dura and cerebellum) in MNI space provided by Shi et al., (2017) using non-linear registration (ANTs SyN); 3) applying the warp file generated in the last step to the ROIs in MNI space with 40-week dHCP T1w template as a reference. Then, we align these ROIs in 40-week dHCP T1w template space with neonate native space using the same procedure as that for the Power atlas. For adults, as the neonatal ROIs in MNI space provided by Shi et al., (2017) have a slightly different spatial resolution (field of view (FOV) = 181 × 217 × 181 mm, spatial resolution of 1 mm isotropic) from the denoised HCP data (FOV = 91 × 109 × 91 mm, spatial resolution of 2 mm isotropic), we first aligning neonatal T1w image (with no dura and cerebellum) in MNI space to MNI T1w template (in 2 mm resolution) using non-linear registration (ANTs SyN) and then applying the warp file generated in the last step to ROIs with MNI T1w template (in resolution 2 mm) as a reference.

#### Traditional algorithms for calculating small worldness

The algorithm for calculating small-world properties proposed by Humphries and Gurney (2008) compares the observed clustering coefficient and characteristic path length with that of equivalent Erdos-Renyi random networks to get the small worldness ($\sigma$) of the observed network, which is given by the following equation

$$\sigma = \frac{\gamma}{\lambda} = \frac{\frac{C_{obs}}{C_{rand}}}{\frac{L_{obs}}{L_{rand}}}, \quad (1)$$

where *C* and *L* stand for clustering coefficient and characteristic path length respectively of the observed network (*obs*) and random network (rand). The equivalent random network has the same number of nodes and the same degree distribution as the observed network. According to this equation, the higher normalized *C* and lower normalized *L*, the higher $\sigma$.

#### Statistical analyses

##### *Comparison of head motion*

We investigated whether there is any significant difference in head motion between the four groups, the adults, full-term neonates, and preterm neonates at or before term-equivalent age (TEA). Independent sample *t*-tests were first applied to test the difference in the mean framewise displacement (FD) between the adults and each neonate group, and between the full-term neonates and each preterm neonate group separately.

##### *Comparison of mean functional connectivity (FC)*

General linear modes (GLMs) were applied to test the difference in mean FC between the adults and each neonate group while controlling the mean FD value, given significantly lower FD in the adults relative to neonates (SI, Figure S1). Independent sample *t*-tests were used to detect the difference in mean FC between full-term neonates and preterm neonates at TEA/before TEA, as no significant difference in FD was observed between neonate groups.

*Comparison of functional connectivity at the nodal level*

Independent-sample *t*-tests were conducted to detect the difference in FC patterns between the full-term neonates and preterm neonates at/before TEA. The FC matrices were standardized to z-scores to allow comparisons across subjects as the full-term neonates had significantly higher mean FC relative to the preterm neonate groups. Correction for the false discovery rate (FDR) was applied to all statistical results at a threshold of $p < 0.05$.

*Comparison of small-world properties at the nodal level*

Independent-sample *t*-tests were conducted to detect the difference in shortest path length patterns between the full-term neonates and preterm neonates at/before TEA. The path length matrices were standardized to z-scores to allow comparisons across subjects as they were calculated directly based on FC matrices. Group-wise comparisons in clustering coefficient at the nodal level were conducted between the full-term neonates and preterm neonates at/before TEA in GLMs including mean FC as a covariate. FDR correction was applied at a threshold of $p < 0.05$.

### Supplementary Results

#### Comparison of head motion

The adults had significantly lower mean FD relative to the full-term neonates ($t$ (407.96) = -9.23, $p < 0.0001$), preterm neonates scanned at TEA ($t$ (80.81) = -4.11, $p < 0.0001$), and preterm neonates scanned before TEA ($t$ (84.82) = -4.80, $p < 0.0001$) (Figure S1). No significant differences were observed in mean FD between neonate groups.

*Figure S1 about here please*

#### Comparison of mean functional connectivity

GLMs were applied to compare the mean FC in the adults and each neonate group while controlling the mean FD. We observed a significant main effect of group for the comparison between the adults and full-term neonates ($F$ (1, 451) = 10.99, $p = 0.001$), which was driven by significantly lower FC in the adults. GLM comparing the adults and preterm neonates at TEA also showed a significant main effect of groups ($F$ (1, 245) = 42.64, $p < 0.0001$), driven by significantly higher mean FC in the adults. Similarly, GLM comparing the adults and preterm neonates before TEA showed a main effect of group ($F$ (1, 243) = 17.66, $p < 0.0001$), again driven by higher mean FC in the adults (Figure S2a).

Independent *t*-tests were applied to compare the mean FC between the full-term neonates and preterm neonates at/before TEA. The full-term neonates had significantly higher mean FC relative to preterm neonates scanned at TEA ($T$ (1, 126.84) = 9.96, $p < 0.0001$) and preterm neonates scanned before TEA ($T$ (1, 346) = 7.11, $p < 0.0001$) (Figure 2b).

*Figure S2 about here please*

#### The effect of premature birth on nodal-level FC and small-world properties

Visual inspection of the connectivity matrices (Figure S3) suggested that each neonate group had less cohesive networks (lower within relative to between-network connectivity) across brain than the adult group. The sensorimotor networks were more adult-like relative to other networks in the neonate groups. The connectivity matrix in the preterm neonates at TEA resembled that in the full-term neonates. Networks in the preterm neonate before TEA were less differentiated/segregated relative to that in neonates at term or TEA.

*Figure S3 about here please*

Full-term neonates had significantly higher within- and between-network FC across the brain relative to preterm neonates before TEA (Figure S4a). By term-equivalent age, the functional connectivity maps in preterm neonates were like those in full-term neonates. However, we still observed significantly different within- and between-network FC between the full-term neonates and preterm neonates at TEA. Specifically, the full-term neonates had significantly higher FC within the default mode network but lower FC within the sensorimotor-hand network relative to the preterm neonates at TEA (Figure S4b). A similar pattern of results was observed when comparing shortest path length between the full-term neonates and preterm neonates (Figure S4).

*Figure S4 about here please*

When comparing nodal-level clustering coefficient, the full-term neonates showed a higher clustering coefficient in the right postcentral gyrus of sensory/somatomotor mouth network and lower clustering coefficient in the right precuneus of default mode network, when compared with preterm neonates before TEA (Figure S5). No significantly different clustering coefficient was observed in any nodes between the full-term neonates and preterm neonates at TEA.

*Figure S5 about here please*

*Reproducibility of results*

*Adults versus neonates* Higher small worldness, higher clustering coefficient and lower path length were consistently observed in the adults relative to preterm neonates before TEA across different algorithms and types of FC matrices (C3 in Figure S6) and parcellations (Figure S7 and Table S2). Using the power atlas, we consistently found significantly higher small worldness and higher clustering coefficient in adults relative to full-term neonates (C1 in Figure S6) and preterm neonates at TEA (C2 in Figure S6) across different algorithms and types of FC matrices. We did not observe any significant difference in small worldness between adults and those two neonate groups while using the UNC atlas (Figure S7a). However, we observed significantly lower path length (Figure S7c) in adults relative to full-term neonates and significantly higher clustering coefficient in adults relative to preterm neonates at TEA (Figure S7b). These findings suggested the adults exhibited a more well-developed small-world architecture relative to neonates.

*Figure S6, S7 and Table S2 about here please*

*Full-term versus preterm neonates* Relative to preterm neonates before TEA, full-term neonates were consistently found to have higher small worldness and lower path length regardless of using different algorithms and types of FC matrices (C4 in Figure S6) and parcellations (Figure S8 and Table S3). In terms of the comparisons between full-term neonates and preterm neonates at TEA, the only subtle difference was in the comparison in small worldness. The marginal decrease in small-world propensity found in preterm neonates at TEA using the new algorithms was found to be significant using the commonly used algorithms (C5 in Figures S6 and Figure S8). Nevertheless, both results suggested a prematurity-related weaker functional small-world architecture in neonates at TEA.

*Figure S8 about here please*

*Preterm neonates at TEA versus before TEA* Lower path length was consistently observed in preterm neonates at TEA relative to the same neonates before TEA regardless of using different algorithms and types of FC matrices (C6 in Figure S6) and parcellations (Figure S9 and Table S3), although some results did not survive multiple comparisons correction. Preterm neonates at TEA showed significantly higher small worldness relative to the same neonates before TEA using the new algorithm across different brain parcellations (Figure S9). However, we did not find any difference in small worldness between the two groups using the traditional algorithm (Figure S6).

*Figure S9 and Table S3 about here please*

### Supplementary Figures

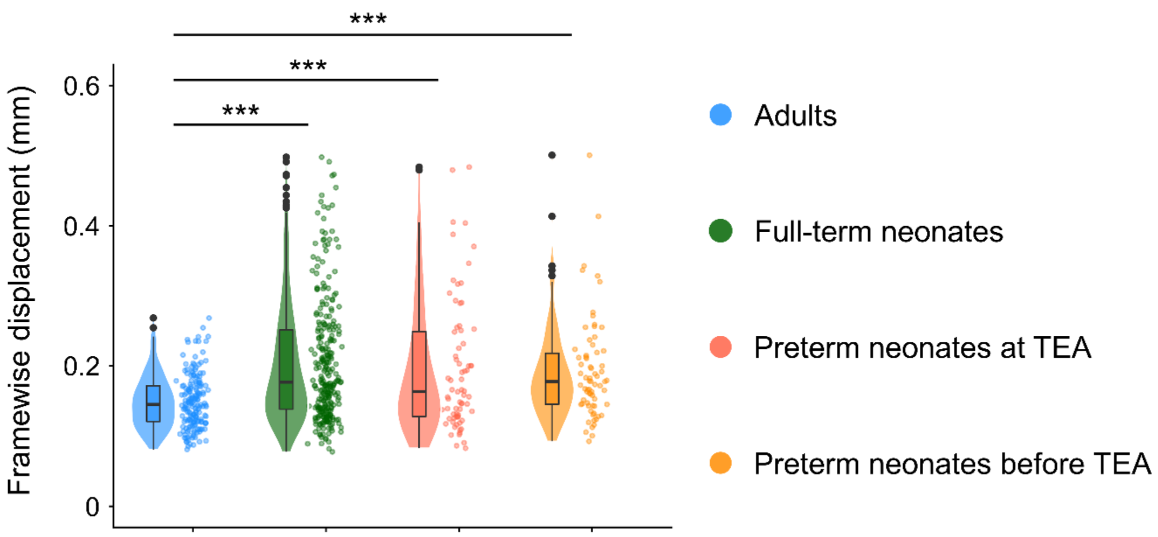


**Figure S1.** Head motion in the adults and neonate groups after censoring. The violin plots show the distribution of the data. The line that divides the box into two parts represents the median of the data. The ends of the box represent the upper (Q3) and lower (Q1) quartiles of the data. The extreme lines of the boxplot show Q3 + 1.5 * interquartile to Q1 – 1.5 * interquartile range. The black dots beyond the extreme lines show potential outliers. Independent-sample *t*-tests were applied to detect the difference between every two groups. Abbreviations: TEA, term-equivalent age. *** = $p < 0.0005$.

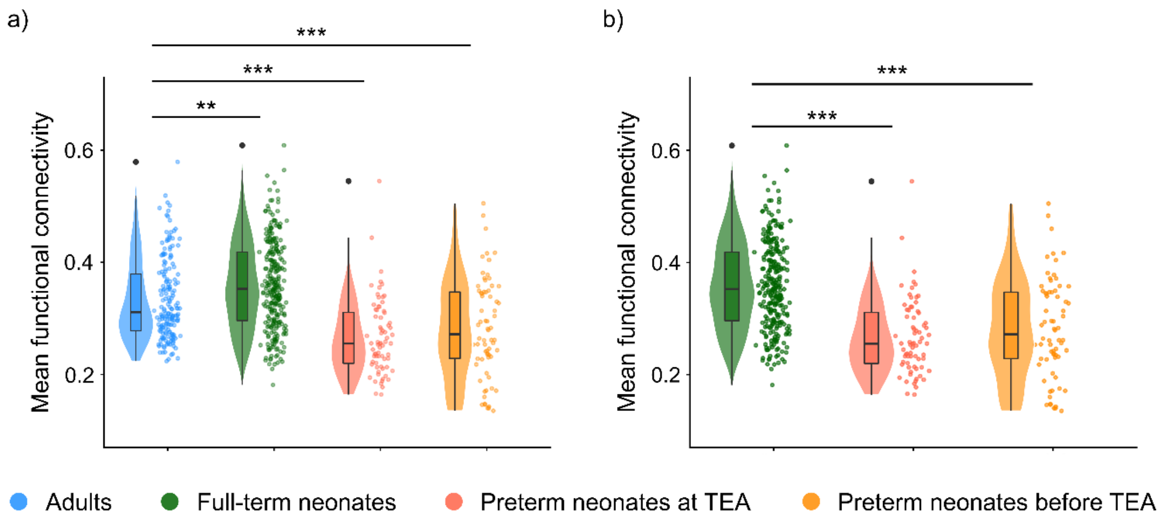


**Figure S2. Mean functional connectivity in adults and neonates.** a) Comparisons between the adults and each neonate group. General linear models were applied to detect the difference in mean functional connectivity (FC) between the adults and each neonate group while controlling head motion. b) Comparisons between the full-term neonates and each preterm group. Independent sample *t*-tests were used to detect the difference in mean FC between full-term neonates and preterm neonates at TEA/before TEA, as no significant difference in FD was observed between neonate groups. The mean FC was calculated for each participant by averaging all positive edges. Abbreviations: TEA, term-equivalent age. *** = $p < 0.0005$; ** = $p < 0.005$.

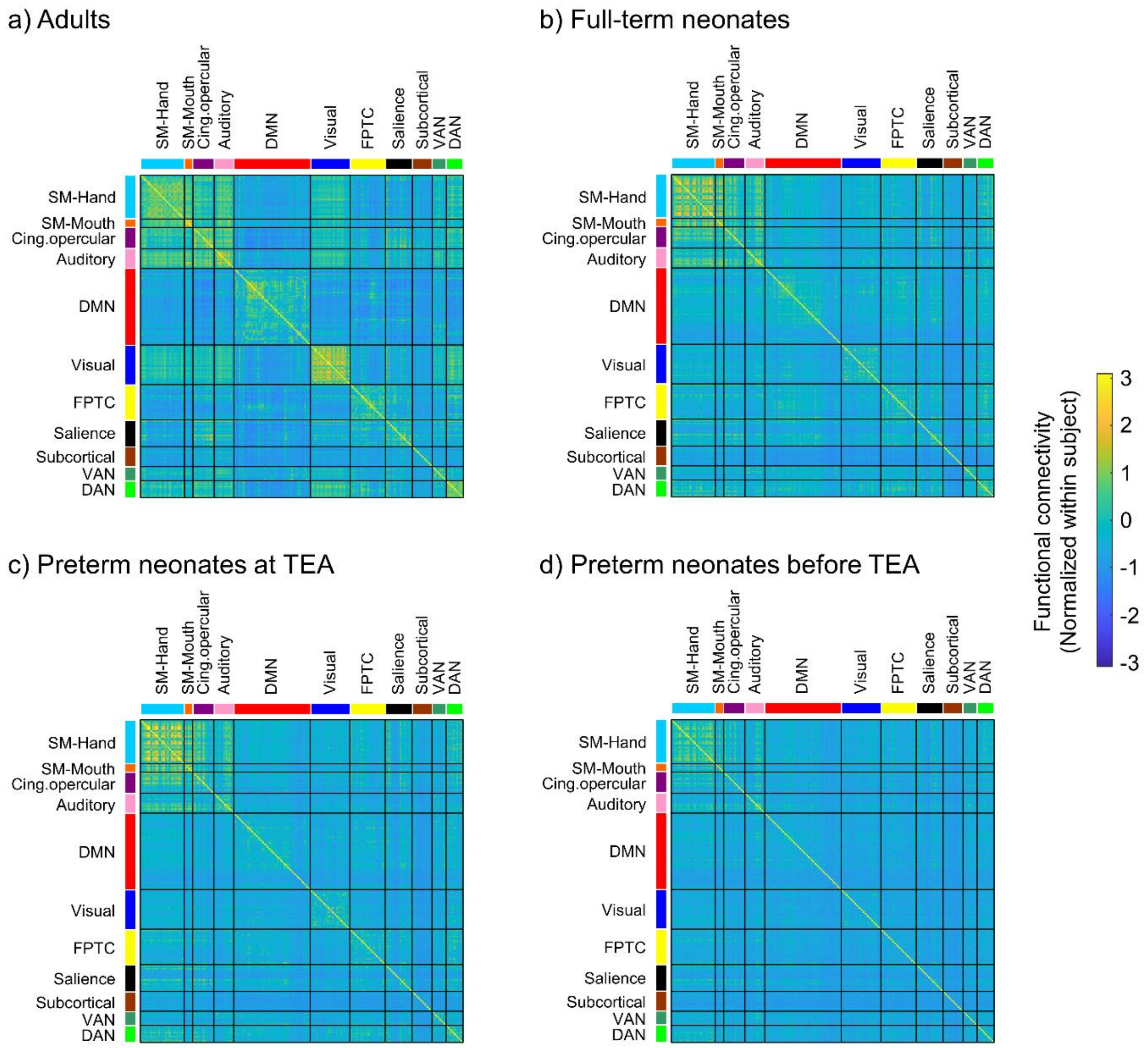


**Figure S3. Functional connectivity in adults and neonate groups.** a) adults, b) full-term neonates, c) preterm neonates scanned at TEA and d) preterm neonates scanned before TEA. The FC value presents here was normalized within each subject before being averaged within each group. Abbreviations: SM-Hand, sensory/somatomotor hand network; SM-Mouth, sensory/somatomotor mouth network; Cing. opercular, cingulo-opercular task control network; DMN, Default mode network; FPTC, frontoparietal task control network; VAN, ventral attention network; DAN, dorsal attention network.

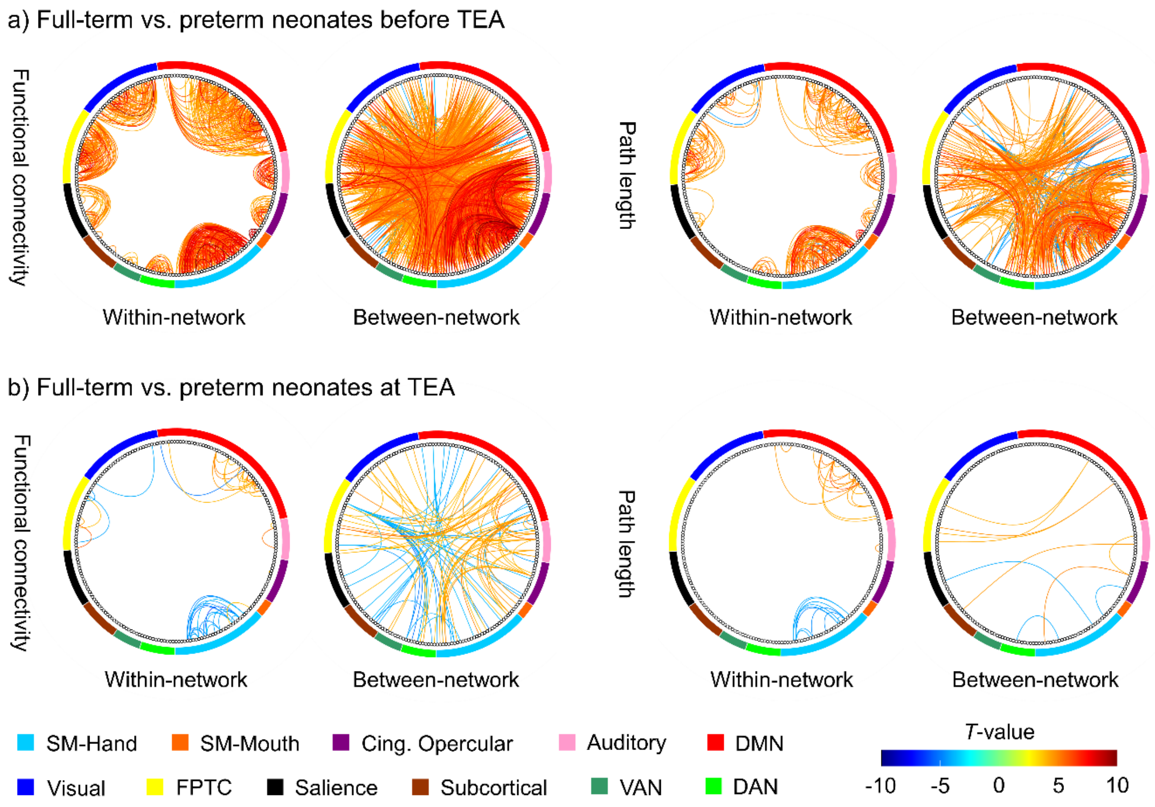


**Figure S4. Differences in functional connectivity and path length between neonate groups.** The lines in warm/cool colour represent higher/lower functional connectivity or shorter/longer path length in the full-term neonates relative to preterm groups. Correction for the false discovery rate was applied at a threshold of $p < 0.05$. Abbreviations: TEA, term-equivalent age; SM-Hand, sensory/somatomotor hand network; SM-Mouth, sensory/somatomotor mouth network; Cing. opercular, cingulo-opercular task control network; DMN, Default mode network; FPTC, frontoparietal task control network; VAN, ventral attention network; DAN, dorsal attention network.

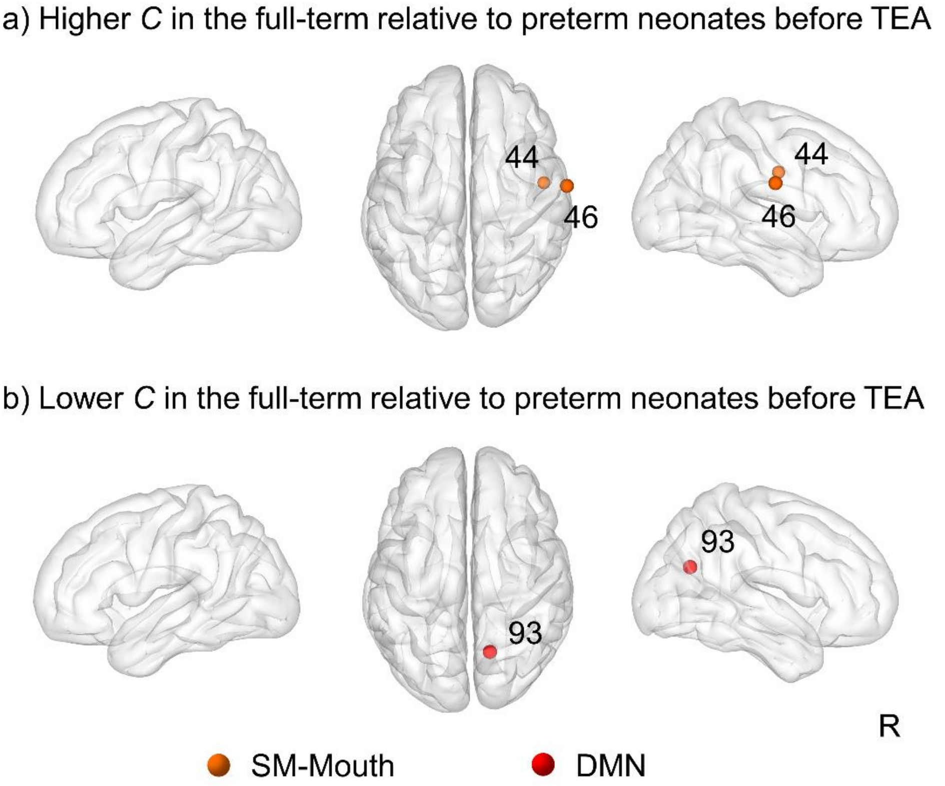


**Figure S5. Effects of premature birth on nodal-level clustering coefficient.** Nodes significantly showing a) higher or b) lower clustering coefficient in the full-term relative to preterm neonates before TEA. Correction for the false discovery rate was applied at a threshold of $p < 0.05$. Abbreviations: TEA, term-equivalent age; SM-Mouth, sensory/somatomotor mouth network; DMN, Default mode network; 44, node 44 (right postcentral gyrus); 46, node 46 (right postcentral gyrus); 93, node 93 (right precuneus); R, right.

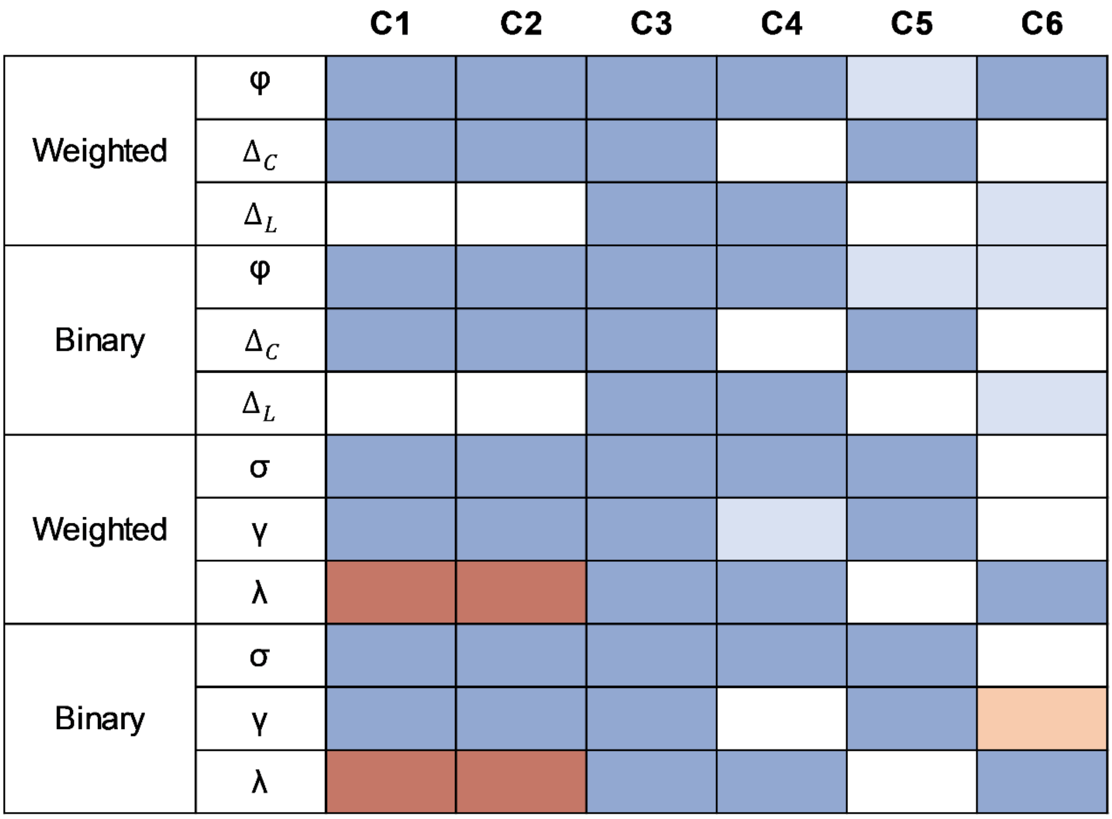


**Figure S6. Consistency of results.** $\varphi/\Delta_C/\Delta_L$ are small-world propensity/normalized clustering coefficient/ characteristic path length calculated according to Muldoon et al., (2016). $\sigma/\gamma/\lambda$ are small-worldness/normalized clustering coefficient/characteristic path length calculated according to Humphries and Gurney (2008). The blue/red colour indicates lower/higher small-worldness ($\varphi/\sigma$), lower/higher clustering coefficient (higher $\Delta_C$/lower $\gamma$), longer/shorter path length ($\Delta_L/\lambda$) in neonate groups relative to adults (C1 ─ C3), preterm neonate groups relative to full-term neonates (C4 and C5), or preterm neonates before TEA relative to preterm neonates at TEA (C6). The darker colour represents the results that survived FDR correction for multiple comparisons, and the lighter colour indicates $p < 0.05$ but did not survive the correction for multiple comparisons. Abbreviations: C1, comparison between the adults and full-term neonates; C2, comparison between the adults and preterm neonates at TEA; C3, comparison between the adults and preterm neonates before TEA; C4, comparison between the full-term neonates and preterm neonates before TEA; C5, comparison between the full-term neonates and preterm neonates at TEA; C6, comparison between preterm neonates scanned at TEA and the same neonates scanned before TEA; Binary, binary functional connectivity matrix; Weighted, weighted functional connectivity matrix.

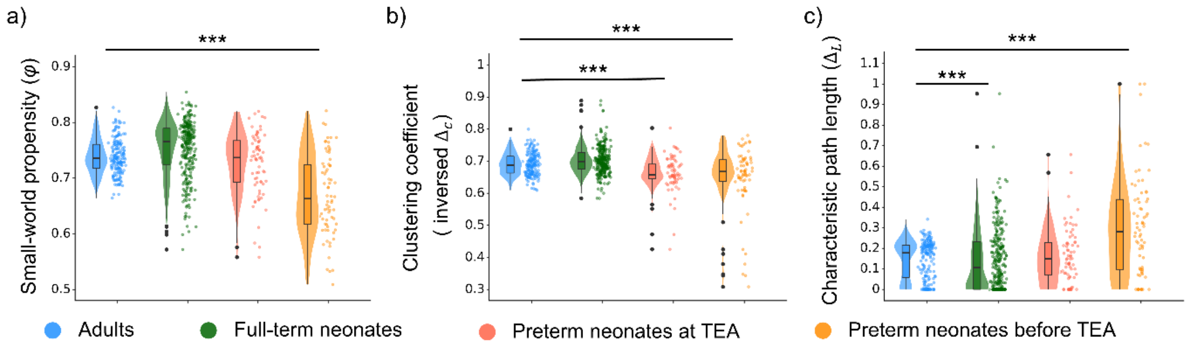


**Figure S7. The development of small-world architecture in the neonates relative to adults (UNC atlas).** a) small-world propensity, b) clustering coefficient, and c) characteristic path length in the neonate groups relative to adults. Higher $\varphi$/inversed $\Delta_C$/$\Delta_L$ represent higher small-world propensity, higher clustering coefficient and longer characteristic path length. The inversed $\Delta_C$ was calculated by 1 - $\Delta_C$ and for visualization purposes only and the statistical analyses were conducted using $\Delta_C$. The violin plots show the distribution of the data. The line that divides the box into two parts represents the median of the data. The ends of the box represent the upper (Q3) and lower (Q1) quartiles of the data. The extreme lines of the boxplot show Q3 + 1.5 * interquartile to Q1 – 1.5 * interquartile range. The black dots beyond the extreme lines show potential outliers. Abbreviations: TEA, term-equivalent age. *** = $p < 0.0005$.

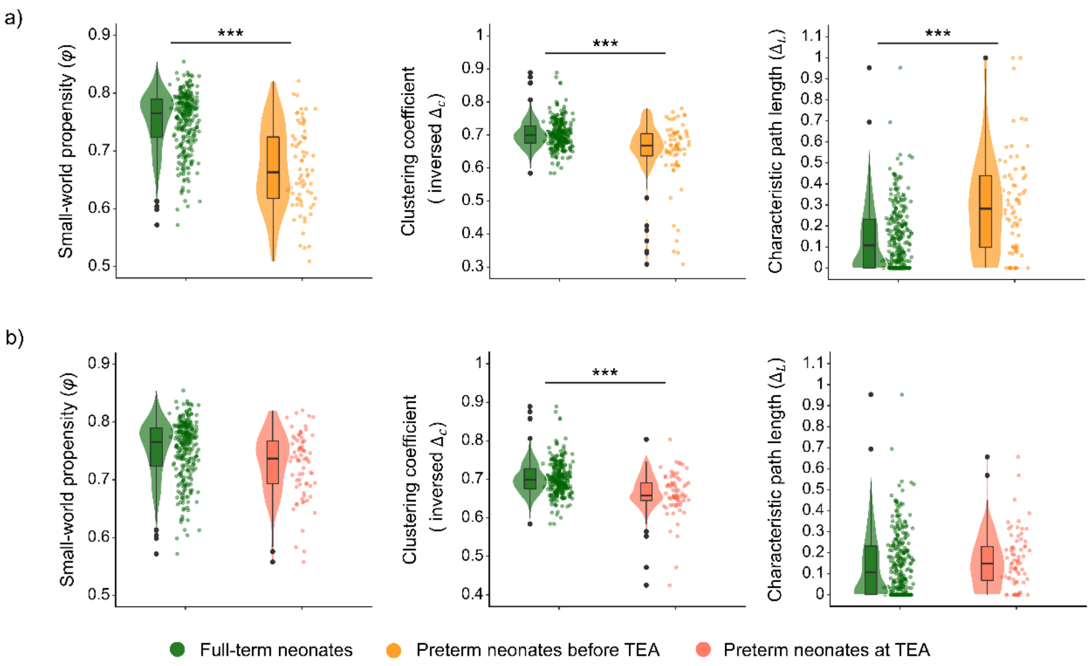


**Figure S8. Effects of premature birth and neonate age on the development of small-world architecture.** a) small-world properties in the full-term neonates relative to preterm neonates before TEA; b) small-world properties in the full-term neonates relative to preterm neonates at TEA. Full-term neonates had significantly higher small-world propensity, higher clustering coefficient and lower normalized path length relative to preterm neonates before TEA, independent of mean functional connectivity. Full-term neonates had significantly higher clustering coefficient relative to preterm neonates at TEA. Higher $\varphi$/inversed $\Delta_C$/$\Delta_L$ represent higher small-world propensity, higher clustering coefficient and longer characteristic path length. The inversed $\Delta_C$ was calculated by 1 - $\Delta_C$ and for visualization purposes only and the statistical analyses were conducted using $\Delta_C$. The violin plots show the distribution of the data. The line that divides the box into two parts represents the median of the data. The ends of the box represent the upper (Q3) and lower (Q1) quartiles of the data. The extreme lines of the boxplot show Q3 + 1.5 * interquartile to Q1 – 1.5 * interquartile range. The black dots beyond the extreme lines show potential outliers. Abbreviations: TEA, term-equivalent age. *** = $p < 0.0005$.

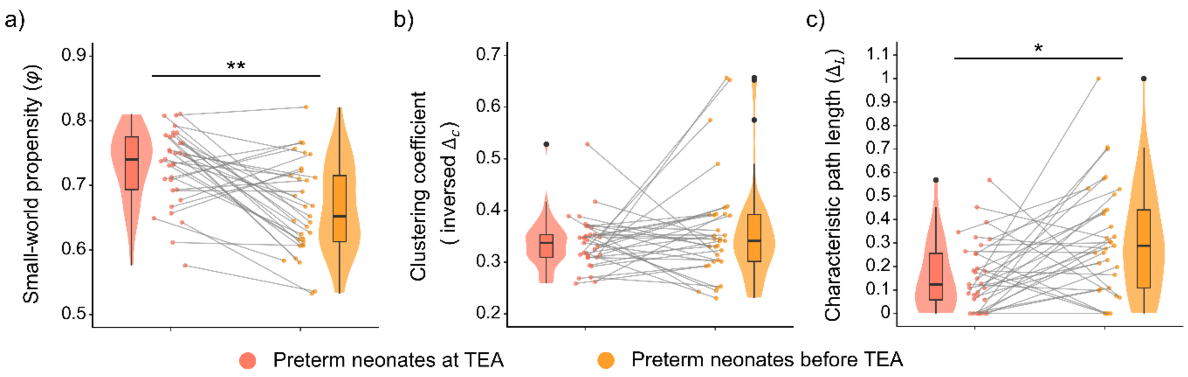


**Figure S9. The development of small-world architecture in preterm neonates up to term-equivalent age (UNC atlas).** a) small-world propensity, b) clustering coefficient, and c) characteristic path length in the preterm neonates scanned at and before term-equivalent age. Significantly higher small-world propensity and lower path length were observed in the preterm neonates at TEA relative to the same neonates scanned before TEA. Higher $\varphi$/inversed $\Delta_C$/$\Delta_L$ represent higher small-world propensity, higher clustering coefficient and longer characteristic path length. The inversed $\Delta_C$ was calculated by 1 - $\Delta_C$ and for visualization purposes only and the statistical analyses were conducted using $\Delta_C$. The violin plots show the distribution of the data. The line that divides the box into two parts represents the median of the data. The ends of the box represent the upper (Q3) and lower (Q1) quartiles of the data. The extreme lines of the boxplot show Q3 + 1.5 * interquartile to Q1 – 1.5 * interquartile range. The black dots beyond the extreme lines show potential outliers. Abbreviations: TEA, term-equivalent age. ** = $p < 0.005$; * = $p < 0.05$.

338

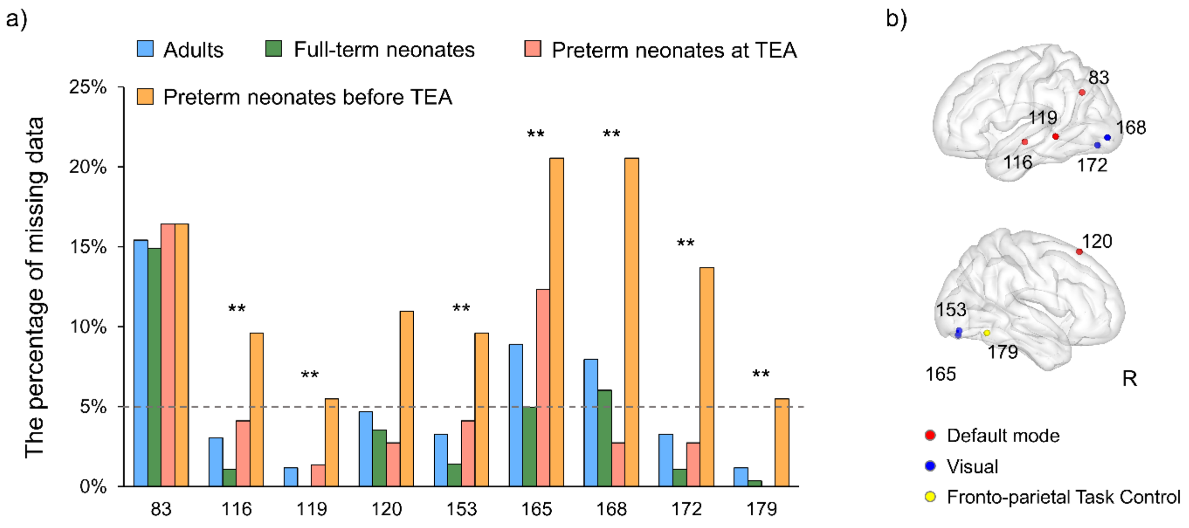


339 **Figure S10. Regions excluded from the Power-264 atlas.** Abbreviations: R, right; ** = $p <$
340 0.005.

**Table S1. Information obtained from the developing Human Connectome Project for scans used in the present study.**

| Group | Sex | Birth age (GA, weeks ± days) | Scan age (PMA, weeks ± days) | Birth weight (Kg) |
|---|---|---|---|---|
| Full-term neonates | 159M/119F | 39.9 ± 8.6 | 41.2 ± 12.2 | 3.35 ± 0.54 |
| Preterm neonates at TEA | 40M/32F | 32.1 ± 25.5 | 40.9 ± 14.6 | 1.77 ± 0.79 |
| Preterm neonates before TEA | 48M/22F | 32.5 ± 20.7 | 34.7 ± 12.7 | 1.79 ± 0.58 |

Abbreviations: GA, gestational age; PMA, postmenstrual age; TEA, term-equivalent age; M, male; F, female.

**Table S2. Comparison in small-world architecture in adults relative to neonates (UNC atlas)**

| **Network properties** | **Comparisons** | ***F*-value (DF)** | ***p*-value** |
|---|---|---|---|
| Small-world propensity | Adults vs. Full-term neonates | 0.58 (1, 450) | 0.447 |
| | Adults vs. preterm neonates at TEA | 2.50 (1, 244) | 0.115 |
| | Adults vs. preterm neonates before TEA | 154.01 (1, 242) | < 0.0001 *** |
| Clustering coefficient | Adults vs. Full-term neonates | 4.63 (1, 450) | 0.032 |
| | Adults vs. preterm neonates at TEA | 18.49 (1, 244) | < 0.0001 *** |
| | Adults vs. preterm neonates before TEA | 30.35 (1, 242) | < 0.0001 *** |
| Characteristic path length | Adults vs. Full-term neonates | 15.28 (1, 450) | < 0.0001 *** |
| | Adults vs. preterm neonates at TEA | 0.73 (1, 244) | 0.394 |
| | Adults vs. preterm neonates before TEA | 49.76 (1, 242) | < 0.0001 *** |

Abbreviations: TEA, term-equivalent age; DF, degree of freedom; ***, $p < 0.0005$.

**Table S3. Comparison in small-world architecture between neonate groups (UNC atlas)**

| **Network properties** | **Comparisons** | ***F*-value (DF)** | ***p*-value** |
|---|---|---|---|
| Small worldness | Full-term vs. preterm neonates at TEA | 2.14 (1, 346) | 0.144 |
| | Full-term vs. preterm neonates before TEA | 124.29 (1, 344) | < 0.001 *** |
| | Preterm neonates at vs. before TEA | 19.52 (1, 31.73) | 0.001 ** |
| Clustering coefficient | Full-term vs. preterm neonates at TEA | 23.48 (1, 346) | < 0.001 *** |
| | Full-term vs. preterm neonates before TEA | 27.65 (1, 344) | < 0.001 *** |
| | Preterm neonates at vs. before TEA | 1.77 (1, 30.85) | 0.193 |
| Characteristic path length | Full-term vs. preterm neonates at TEA | 2.83 (1, 346) | 0.093 |
| | Full-term vs. preterm neonates before TEA | 41.29 (1, 344) | < 0.001 *** |
| | Preterm neonates at vs. before TEA | 7.15 (1, 31.95) | 0.012 * |

Abbreviations: TEA, term-equivalent age; DF, degree of freedom; ***, $p < 0.0005$; **, $p < 0.005$; *, $p < 0.05$.

### Supplementary References


Fitzgibbon, S. P., Harrison, S. J., Jenkinson, M., Baxter, L., Robinson, E. C., Bastiani, M., ... & Andersson, J. (2020). The developing Human Connectome Project (dHCP) automated resting-state functional processing framework for newborn infants. *Neuroimage*, 223, 117303.

Glasser, M. F., Sotiropoulos, S. N., Wilson, J. A., Coalson, T. S., Fischl, B., Andersson, J. L., ... & Wu-Minn HCP Consortium. (2013). The minimal preprocessing pipelines for the Human Connectome Project. *Neuroimage*, 80, 105-124.

Humphries, M. D., & Gurney, K. (2008). Network 'small-world-ness': a quantitative method for determining canonical network equivalence. *PloS one*, 3(4), e0002051.

Power, J. D., Cohen, A. L., Nelson, S. M., Wig, G. S., Barnes, K. A., Church, J. A., ... & Petersen, S. E. (2011). Functional network organization of the human brain. *Neuron*, 72(4), 665-678.

Schuh, A., Makropoulos, A., Robinson, E. C., Cordero-Grande, L., Hughes, E., Hutter, J., ... & Rueckert, D. (2018). Unbiased construction of a temporally consistent morphological atlas of neonatal brain development. *BioRxiv*, 251512.

Shi, F., Salzwedel, A. P., Lin, W., Gilmore, J. H., & Gao, W. (2018). Functional brain parcellations of the infant brain and the associated developmental trends. *Cerebral Cortex*, 28(4), 1358-1368.

Shi, F., Yap, P. T., Wu, G., Jia, H., Gilmore, J. H., Lin, W., & Shen, D. (2011). Infant brain atlases from neonates to 1-and 2-year-olds. *PloS one*, 6(4), e18746.
